# Faithful Scanning Electron Microscopic (SEM) visualization of 3D printed alginate-based scaffolds

**DOI:** 10.1101/2020.03.18.997668

**Authors:** Marcus Koch, Małgorzata K. Włodarczyk-Biegun

## Abstract

The morphological characterization of 3D printed hydrogel-based scaffolds is essential for monitoring their size, shape, surface texture and internal structure. Among other microscopic techniques, Scanning Electron Microscopy (SEM) is capable of visualizing nearly all kinds of materials at different length scales, with exceptional precision, if investigation under vacuum is possible. However, due to the high water content of hydrogel-based scaffolds and the connected volume change after drying, special preparation techniques are necessary to stabilize the 3D architecture when imaged by SEM. Here we present a straightforward cryo-SEM technique to visualize 3D printed hydrogel-based alginate scaffolds. By use of a homemade cryo-SEM holder and plunge-freezing in liquid ethane, scaffolds are visualized from the top and cross-sectional view at different magnifications. The proposed method is compared with SEM imaging in different modes (cyro-SEM, conventional SEM, ESEM) following other commonly used sample preparation techniques, such as plunging in liquid nitrogen, air-drying, freeze-drying and plunging in liquid ethane after graded dehydration. These approaches, except ESEM, lead to shrinkage, deformation, distortion or disintegration of the scaffolds and consequently give rise to artifacts in imaging. The presented results indicate that cryo-SEM after plunging in liquid ethane allows for the most faithful and time-efficient visualization of 3D printed alginate scaffolds.

## 1 Introduction

3D bioprinting is a powerful fabrication technique for use in tissue engineering (TE) due to the possibility of obtaining complex, predesigned structures with precise control over material deposition, down to several micrometers [1, 2]. One of the most widely investigated approaches is the extrusion printing of hydrogel-based materials (so-called hydrogel inks) [3]. In recent years much attention was devoted to the development of functional inks, yet, new well-printable, biocompatible inks are still needed for TE applications [4]. To characterize and compare printability and quality of the developed inks, several protocols were proposed, mostly based on the observation and quantification of the shape of the printed strands and scaffolds [3, 5-11]. For the detailed ink characterization, the proper visualization of the printed structures, regarding their morphology, microstructure and surface characteristics, plays an important role.

To visualize printed structures, the use of bright field or stereo light microscopy (LM) [3, 5-8, 10, 12, 13], fluorescent or confocal microscopy [13-18] and scanning electron microscopy (SEM) [18-26] was reported. For transparent hydrogel-based inks, light microscopy is limited by the weak scattering of the light, leading to the low contrast even under optimized illumination (e.g. dark-field or oblique). A possible solution is dye addition to the ink to colorize printed structure and increase the contrast [3, 5-7, 12]. However, this requires additional preparation steps, interference with the ink composition and the resolution of the analysis is still limited, by Abbe’s law, to λ/2 or 200 nm [27]. Fluorescent and confocal imaging usually offer information at a higher level of detail, but often require more complex preparation procedures (fluorescently-labeled ink preparation or post-printing scaffold staining); they are also limited in time due to the photo-bleaching of the dye, and may lead to the photo-damage of the investigated sample.

Scanning electron microscopy (SEM) can serve as a powerful, complementary tool for optical imaging of the printed structures. It allows for detailed morphological analysis, from scaffolds overall architecture to strands surface, cross-section and material microstructure, due to the high resolution (down to the nanometer range), high depth of field, and a vast range of available magnifications, commonly from 10 x to 500.000 x [28]. However, because of the high vacuum conditions required for imaging, analyzed samples typically need to be in a dry solid state for investigations [28]. Therefore, high water content of the hydrogel-based inks complicates their proper structural characterization by SEM technique [29]. Sample preparation methods must preserve the original scaffolds structure, without leading to artifacts or lost information.

The most common approaches of hydrogel-based printed scaffolds preparation for SEM can be divided into two groups: (1) dehydration using a gradation series of ethanol, followed by additional drying step: air drying [12, 24, 30], drying in hexamethyldisilazane [31, 32], critical point drying [18, 21, 23] or freeze-drying [14]; (2) freeze-drying of the fresh, water-containing samples [4, 19, 22, 25]. Prior to the dehydration procedure, the samples containing biological species (e.g. cells) are usually fixed with glutaraldehyde [18, 23, 24, 30, 31]. The dry samples can be additionally coated with a metal (e.g. gold or platinum) or carbon layer before examination [14, 18, 23, 31] to avoid sample charging and increase contrast when imaging under high vacuum conditions.

These preparation techniques bear the risk of irreversible modifications of the sample, which can lead to imaging artifacts. Hydrogel dehydration using ethanol series can lead to shrinkage and densification of the sample [33, 34]. Freeze-drying or critical point drying of hydrogels leads to formation of porous network of the dehydrated samples [29, 35-41]. During the first freezing, the ice crystals precipitate from the single-phase mixture of polymer compounds and water, and the resulting hydrogel phase decreases in water content [42-46]. Under exposure to vacuum, the ice crystals sublimate, typically at temperatures above 173 K/-100°C [47], resulting in a porous polymer network [39, 48]. As a result, an imaged structure does not reflect the original hydrogel state [29, 44, 48].

To investigate hydrogel samples without essential water loss, Environmental Scanning Electron Microscopy (ESEM) [49, 50] and cryo-SEM [29, 51] can be used. However, one of the limitations of ESEM technique is a restricted field of view (to our knowledge to 500 µm [52-54]), so only small areas of printed scaffolds can be visualized in the wet/humid state or image stitching needs to be applied. In contrast, cryo-SEM is capable of visualizing the same field of view as conventional SEM. Dependent on the working distance, the field of view can exceed 10 mm × 10 mm. Ensikat et al. have proposed an approach allowing for fast and straightforward sample preparation for cryo-SEM by the use of a passive cooling holder without external connections for continuous cooling [55]. The sample is fixed on the holder (metal block) and slowly (over around 5 minutes) cooled in the liquid nitrogen to 77 K/-196°C. The holder with the sample is directly moved to SEM and used for imaging. Using this technique, different kinds of biological samples, like plants, bacteria or fungi, were successfully investigated in high vacuum conditions for several hours at temperatures below 193 K/-80°C [55]. During longer investigations, the warming-up can lead to increased freeze-drying and sublimation of water [56], causing structural changes. If the SEM is equipped with a turbo-molecular pump, imaging of the sample can start at temperatures near T = 77 K/-196°C within several minutes, allowing shorter investigation times than for conventional cryo-SEM. Additionally, the intrinsic electrical conductivity of ice is high enough to investigate the frozen samples without any conductive coating, as is often required for SEM, at low accelerating voltages (up to 5 kV) and small beam intensities (up to 20 pA) [29, 55].

To prepare wet samples for cryo-SEM, plunge-freezing [57-59] was reported for the investigation of protein structures [60], liposomes [61] and nanoparticle solutions [62]. Following extremely fast cooling (>10.000 K/s), wet samples can be preserved in their nearly natural surroundings and artifacts due to the drying of the samples are largely prevented [57]. Plunging is typically done into an undercooled liquid, like ethane or propane at T < 110 K/-163°C, because the freezing velocity is higher than in plunging into liquid nitrogen [57]. The state of ice (amorphous or crystalline) after plunge-freezing depends on the freezing velocity, the ambient pressure during plunging, and the thickness and the composition of the sample [57]. The amorphous state is highly desirable as crystallization of ice can lead to a structural change in the sample due to the mechanical strain associated with the phase separation. Wet samples up to several µm thickness can be plunged with the formation of amorphous ice at atmospheric pressure [29, 57]. For thicker samples, a high-pressure freezer is necessary or cryo-protectants need to be added to preserve the amorphous character of the wet samples [57, 58]. However, as printed scaffolds are typically much thicker than several µm, their size is not suitable for high-pressure freezing. Plunging of scaffolds at atmospheric pressure in undercooled liquid ethane is expected to lead to the precipitation of small ice crystals in the micrometer range inside the polymeric network; these ice crystals will sublimate during the investigation in the vacuum of the SEM chamber resulting in a porous structure. Therefore, unaffected imaging after plunging is a function of imaging time and magnification [39].

SEM investigations of hydrogels, including a detailed discussion regarding the challenges of preventing artifacts formation, can be found e.g. for alginate bulk hydrogels [29, 35, 63], carrageenan beads [35] or hydrated bacteria biofilms [28, 64]. Such detailed investigations are missing for 3D printed hydrogel-based scaffolds. The topic appears to be highly relevant as SEM imaging is commonly used in scientific reports on the development of new inks and on scaffold printing [11, 18-24, 26].

The aim of this study was to print alginate-based inks and compare different, commonly applied techniques of hydrogel sample preparation for subsequent SEM investigation. We present a detailed protocol, developed in our lab, for plunging in liquid ethane followed by cryo-SEM. It is argued that this technique has several advantages over other preparation methods: besides being fast and easily adaptable for any types of printed hydrogels, microstructures can be imaged faithfully and essentially artifact-free, which is a necessary prerequisite for developing new inks and optimizing 3D printing of such materials.

## 2 Materials and Methods

### 2.1 Materials

Chitosan (Ch), in the form of high molecular weight powder, was obtained from Fluka BioChemika (Switzerland); alginic acid sodium salt, high viscosity, was obtained from Alfa Aesar (Germany). All other chemicals were purchased from Sigma-Aldrich (Germany). Printing needles were provided by Vieweg (Germany), Optimum® cartridges by Nordson (Germany).

### 2.2 Ink and bath solution preparation

Inks composed of 4% (w/v) solution of alginic acid sodium salt (medium viscosity) in Milli-Q water were prepared. The samples were physically mixed with a spatula, followed by the use of NeoLab multifunctional mixer (few hours or overnight) until homogeneous solutions were obtained. Solutions were loaded into 3 ml or 10 ml printing cartridges.

Bath solutions were prepared by mixing in 1:1 volume ratio: 0.05% (w/v) chitosan in 2% (v/v) acetic acid solution with 10 mM CaCl_2_, leading to 0.05% (w/v) chitosan and 5 mM CaCl_2_ in the final printing bath.

### 2.3 Scaffolds printing

Alginate-based 2-layers grid-like constructs were printed using square design with ca. 1 cm x 1cm dimensions (1.5 mm strand distance), and perpendicular layers placement; using 20 – 40 kPa pressure and 10 mm/s printing speed. Printing was performed directly into the bath solution, on the glass slides fixed with type stripes in 6 well-plates or directly into the plate, using 250 µm inner diameter conical needle (gauge 25), with cartridge temperature T = 313 K/40°C and bath temperature T ∼ 308 K/35°C. Z-offset of the first layer was ca. 150 µm and layer height was set to 180 µm to ensure a good connection between glass and consecutive printed layers. Printers BioScaffolder 3.2 (GeSiM, Germany) and Bio X (Cellink, Sweden) were used. Prior to imaging, samples were stored in 100 mM CaCl_2_ solution at T = 277 K/4°C for at least 2 hours, and up to 11 days, for increasing scaffolds stability. No substantial changes of the scaffold structure in this storage time could be observed (see Supplementary Information, Figure S1).

### 2.4 Light microscopic imaging

The freshly prepared or stored (up to 11 days) printed scaffolds were imaged by stereomicroscopy under opaque illumination (Olympus SZX 16 equipped with an Olympus SC50 CCD camera). Images were acquired to characterize the as-prepared state after printing/storing in solution.

### 2.5 Cryo-SEM: sample preparation and imaging

The same samples were used for light microscopy and SEM imaging investigation. The time between light microscopy characterization and cryo-SEM sample preparation was in the range of 10 – 15 minutes. The size of the plastic or glass support holding the scaffold was adjusted in x-y dimension to approximately 12 mm × 20 mm using a ruler (for glass slides) or pincers (for plastic well plate), in order to fit the size of the plunger. The samples were washed once with demineralized water and then the excess of water was carefully removed with soft tissue.

A Gatan (Pleasonton, Or, USA) CP3 cryo plunger was used to shock-freeze the samples in liquid ethane at T = 108 K/-165°C. Next, an own-built cryo-SEM brass holder (see Figure 1 A) was placed into liquid nitrogen and cooled down to T = 77 K/-196°C (Figure 1 B). A Teflon cup was placed under the metallic sample holder block to reduce heat transfer to the SEM stage. Temperature measurements of the cryo-SEM holder were performed using a Pt100 thermo-couple. The samples were transferred from the plunge freezer to the cryo-SEM holder under liquid nitrogen and placed in the center, under a tilting angle of 50° (Figure 1 C). For cross-section analysis, freeze-fracture was done at this stage using a pre-cooled knife. In the last step, the cryo-SEM holder was completely covered with liquid nitrogen prior to vacuum initialization, in order to avoid ice crystal formation on the sample caused by air humidity. The time between the initialization of vacuum in the chamber and the start of the SEM investigation was in the range of 2 – 3 minutes. Due to the evaporation of nitrogen, the Joule-Thompson effect was observed: a decrease of the temperature to T = 71 K/-202°C was followed by a slow rise to T = 150 K/-123°C within 45 min of vacuum (Figure 1 D). Without the use of the Teflon cup, heat transfer to the SEM sample stage dominated the first increase in temperature and the resulting temperature curve was shifted by approximately 50 K/°C to higher temperatures (Figure 1 D).

**Figure 1.**
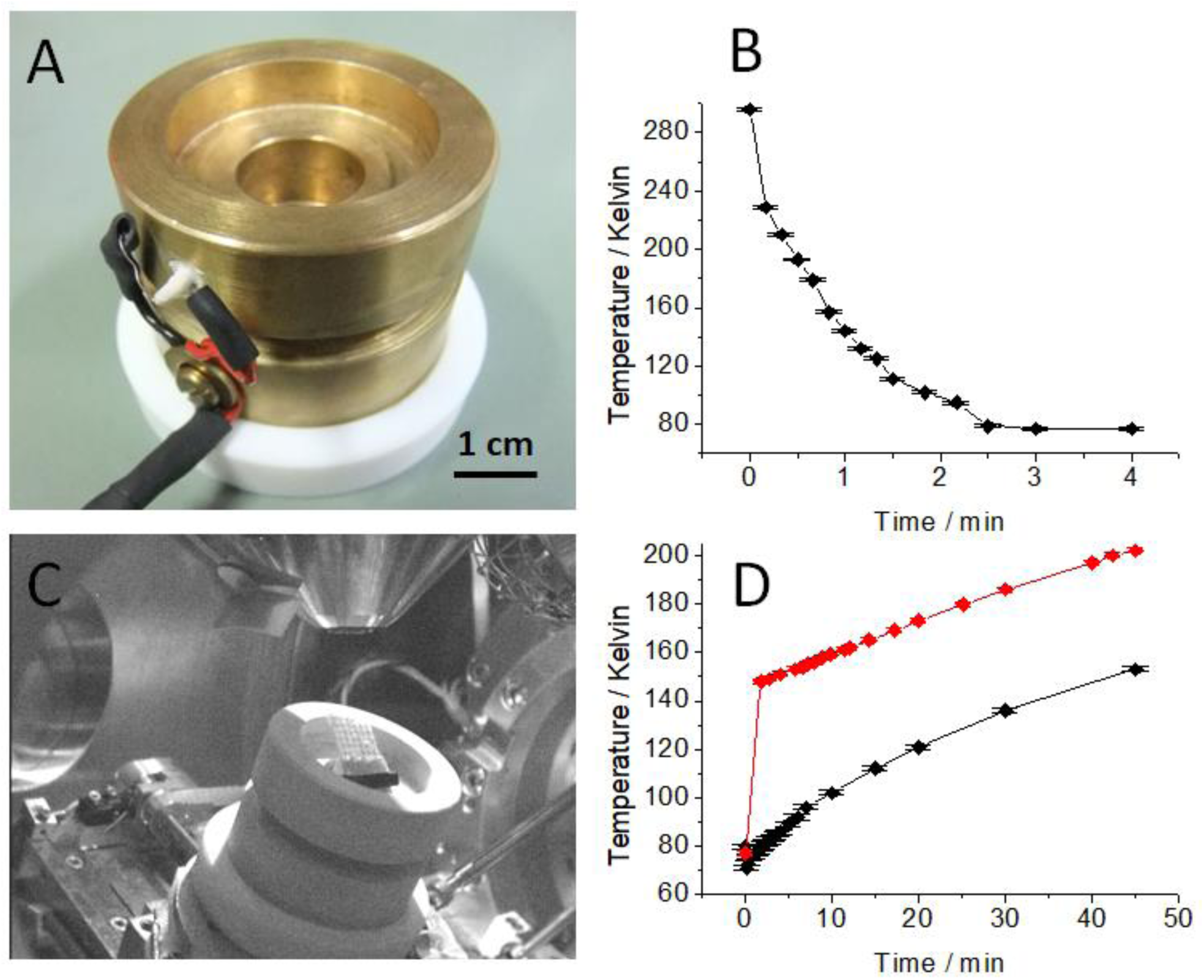
Cryo-SEM investigation setup: Passive home-made cryo-SEM sample holder with Pt100 thermo-couple and Teflon cup to reduce heat transfer to the SEM stage (A); Temperature change of the sample holder after placing in liquid nitrogen (B); Tilted cryo-SEM sample holder inside the SEM chamber (C); Temperature evolution of the cryo-SEM holder inside the SEM chamber after vacuum initialization with Teflon cup (black curve) and without Teflon cup (red curve) (D).

For plunge-freezing in liquid nitrogen the samples were immersed directly into the liquid nitrogen bath of the cryo-SEM holder and mounted under a tilting angle of 50°.

To analyze the changes in the structure of the hydrogel after drying, dehydration with ethanol series was done using ethanol/water mixtures with increasing ethanol concentration. Samples were incubated in consecutive steps for 10 min in 30%, 50%, 70%, 80%, 90% and 96% ethanol/water mixtures and twice for 20 min in pure ethanol. Excess ethanol was carefully removed by a soft tissue, followed by plunging in liquid ethane and cryo-SEM investigation as described above.

An FEI (Thermo Scientific, Dreiech, Germany) Quanta 400 FEG equipped with a turbo-molecular pump was used. Secondary electron images were acquired in high vacuum mode at 3 or 5 kV accelerating voltage with a spot size of 3 and a dwell time of 3 µs. In the cases when water vapor from the atmosphere had led to ice crystals formation on top of the surface during sample transfer, samples were investigated after sublimation of these ice crystals inside the SEM chamber. By using the tilting option of the sample stage, the scaffolds were investigated under different angles. A field of view up to 10 mm × 10 mm was achieved. Magnifications up to 5.000x were obtained with good resolution without a need for a conductive coating of the sample (see Figure S7). After SEM imaging the samples were discarded and the cryo-holder was kept under liquid nitrogen for the next run.

### 2.6 SEM: sample preparation and imaging

All samples were washed with demineralized water and excess of solvent was carefully removed with soft tissue. Three different sample preparation procedures were performed as follows. (1) Samples were air-dried under ambient conditions overnight. Then they were placed directly onto the stage of the SEM and imaged under high and low vacuum conditions at T = 293 K/20°C. (2) To simulate freeze-drying, the samples were stored (in a box filled with water droplets to maintain equilibrium conditions) in the freezer at T = 253 K/-20°C overnight. Samples were directly placed onto the stage of the SEM and investigated inside the SEM under high vacuum during thawing to T = 293 K/20°C. After thawing, the dried samples were imaged under low vacuum conditions. (3) Wet samples were placed on the SEM stage at room temperature and a vacuum was initialized, leading to sample drying inside the SEM. Imaging was performed, both in high and low vacuum, when no further change in vacuum conditions was observed.

Imaging was performed using an FEI Quanta 400 FEG, the same equipment as used for cryo-SEM, at 5, 10 and 20 kV accelerating voltage. Dependent on the conductivity of the sample and the resulting charging using high vacuum conditions during imaging, dried samples were also investigated in low vacuum mode (p = 100 Pa water vapor).

### 2.7 ESEM: sample preparation and imaging

After washing with demineralized water, the samples were directly placed onto an aluminum sample holder using heat conductive carbon tape and investigated in ESEM mode starting at p = 800 Pa after cooling the sample to T = 276 K/3°C. Under these conditions the wet state of the samples is preserved at the beginning of the experiment.

To continuously observe dynamic changes in the structure of the printed hydrogel strand during the dehydration process, in-situ freeze-drying was performed using the ESEM mode starting at p = 800 Pa and T = 276 K/3°C by cooling the hydrogel sample stepwise to T = 253 K/-20°C while reducing the pressure stepwise to 100 Pa (according to the p-T phase diagram of water), followed by thawing to T = 276 K/3°C at 100 Pa. Images were acquired continuously every 15 to 90 seconds.

Imaging was performed with FEI Quanta 400 FEG, the same equipment as used for cryo-SEM, at 10 kV accelerating voltage using the ESEM mode together with the Peltier cooling stage from FEI.

## 3 Results and Discussion

### 3.1 Scaffolds printing

Alginate (medium or high viscosity, 4% (w/v) solution) was chosen for this study as an exemplary hydrogel-based printable material due to its wide use for bioprinting [65, 66]. In order to maintain the designed shape, the material was printed directly into the bath solution containing a mixture of CaCl_2_ for ionic crosslinking and chitosan at pH ∼3. The chitosan (polycation) interacts with alginate (polyanion) to form a skin layer of the polyion complex on the printed strands. Scaffolds with well-defined strands were obtained. Merging and collapse of the layers due to gravity, commonly mentioned as a drawback of alginate printing [19], were limited (see Figure 2B and, for more information, Supplementary Information).

**Figure 2.**
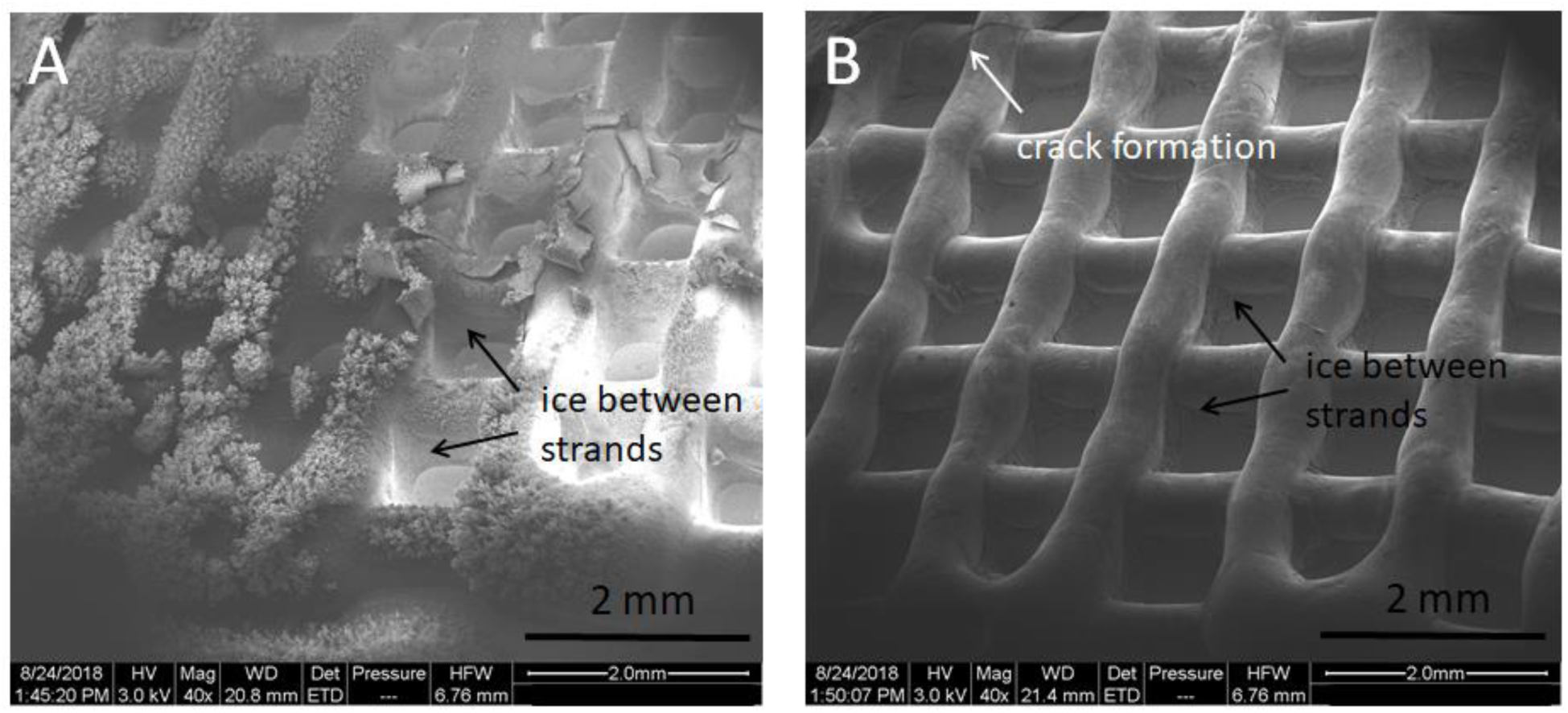
3D printed scaffold with ice crystals on top and between the strands directly after plunging in liquid ethane and imaging at T = 80 K/-193°C (A) and after sublimation of ice at T = 95 K/-178°C after 8 minutes in a vacuum (B). As ice sublimation progresses, the native structure is revealed, followed by crack formation at longer times.

### 3.2 Imaging of printed scaffolds with cryo-SEM

Plunging in liquid ethane or plunging in liquid nitrogen of hydrated samples was applied prior to cryo-SEM visualization. By plunge-freezing, the use of a home-made sample holder and a quick transfer of the sample to the cryo-SEM holder under liquid nitrogen, aimed at maintaining a shock-frozen state of the hydrogel with reduced ice crystal formation. The comparison between LM and SEM top-view images of the scaffolds (Figure 3) reveals well maintained initial shape after applying both preparation techniques. However, the ice formation was not fully avoided (Figure 2). The ice crystals visible on the top of the scaffolds originated from the unbound water, present in different amounts, depending on the 3D architecture of the scaffolds and the efficiency of the drying with a tissue prior to plunging. Additional formation of ice on the scaffold surface was caused by vapor condensation from the atmosphere. Applying a vacuum inside the SEM chamber led to ice sublimation. Several minutes after vacuum initialization the ice crystals formed on the top and between the strands were not visible anymore (Figure 2), revealing the native structure of the printed hydrogels. The scaffolds were imaged with very little further morphological change for several minutes up to one hour (see Figure S2 in Supplementary Information). This time is dependent on the scaffold geometry, printing material and substrate type, as all these factors influence the process of thawing. At longer times, shrinkage of the strands, formation of the elongated structures in contact with the substrate due to the stretching of the adherent material, crack formation and a roughening of the surface were observed (see Supplementary Information and Figure S3A-B). This observation can be explained by the subsequent sublimation of the ice formed inside the strands and originating from the unbound and bound water of the hydrogel. The prolonged sublimation led to the freeze-drying effect [29]. The polymeric structure collapsed and the strain on the material increased, which ultimately caused crack formation to relieve the resulting stress. As the mproposed home-made cryo-SEM holder was a metal block equipped with the Teflon cup, the increase in sample temperature associated with the sublimation of ice crystals from the sample surface in a vacuum was slower (compared to the cryo-SEM holder without Teflon cup). As a result, the proposed setup allows for longer times of imaging with minimized artifacts formation.

For both preparation techniques, the 3D structure of cryo-SEM imaged samples and surface morphology were well preserved with little evidence of major structural rearrangement, provided the sublimation time was not too long. The architecture of the individual strands and separation between printed layers were clearly visible, as shown in the top-view and titled-view images (Figure 2 and Figure 3; see as well Supplementary Information for more images). The surface of the strand was non-porous, with detectable rims that indicated the direction of material flow during or after printing (Figure S3A-B, S7, S14). At higher magnification, a slight shrinkage, visible as parallel wrinkles, could be noticed (Figure S3A-B). Although the freezing velocity using undercooled liquid ethane was higher compared to liquid nitrogen, no signs of the differences between preparation with liquid ethane and nitrogen were observed.

**Figure 3.**
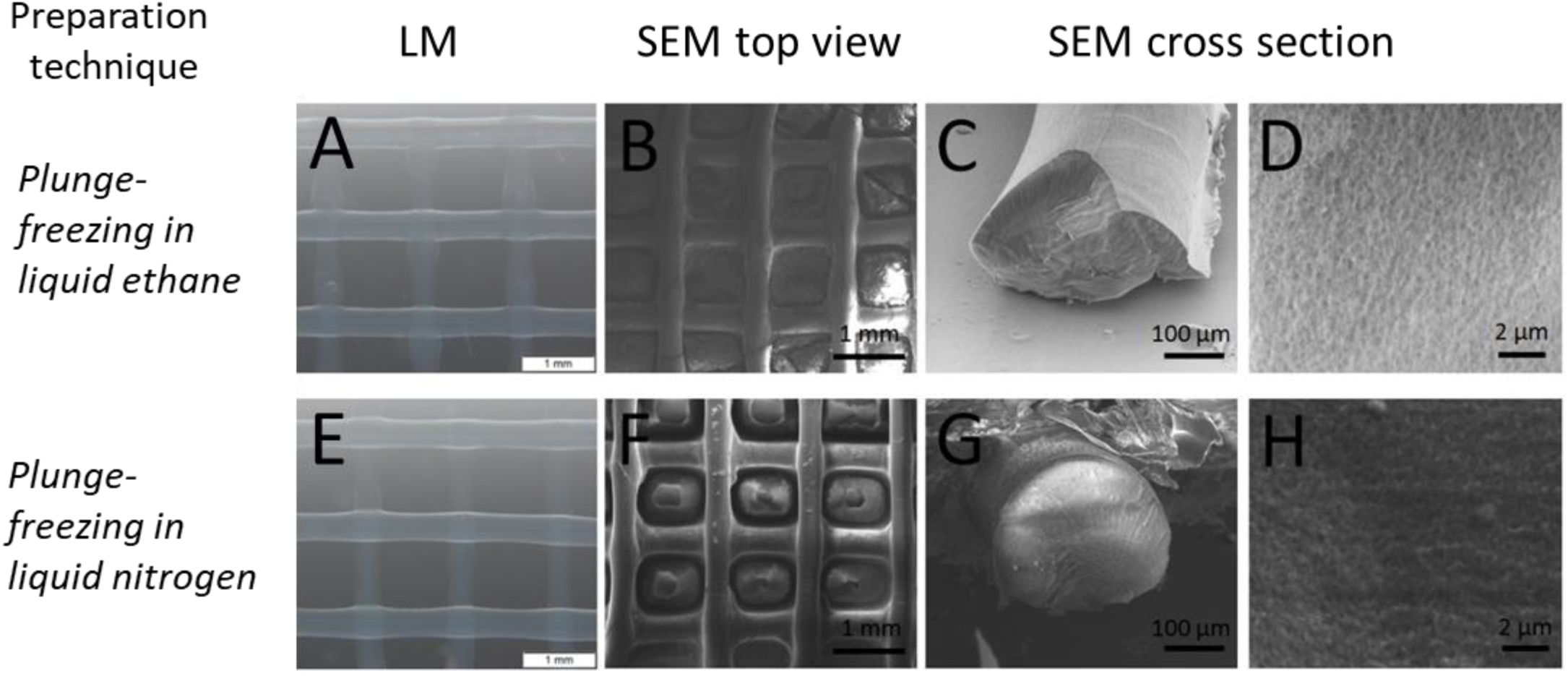
Cryo-SEM investigation: Light microscopic images of 3D printed scaffolds as received (A, E). SEM images after plunge-freezing in liquid ethane (B: top view; C-D: cross-section) and plunge-freezing in liquid nitrogen (F: top view; G-H: cross-section). Scaffolds 3D architecture is well preserved, artificial porosity inside the stands is visible at higher magnification.

Cross-sections of the scaffolds, visible after freeze-fracture (Figure 3), revealed the well-preserved round-shaped profile of the strands. The view allowed observation of the internal structure of the hydrogel: at lower magnification a dense structure was observed (Figure 3C, G and Figure S14M), while at high magnifications (original mag = 20.000x) pores in the sub-micrometer range were noticed (Figure 3D, H). The porosity results from the ice sublimation in the vacuum after precipitation of ice during solidification of the hydrogel, as described above, and is not an inherent property of the printed material. For the samples with small volumes, the ice crystallization for the high freezing velocities typical for liquid ethane or liquid nitrogen at the sample surface was minimized. However, due to the relatively high thickness of the printed strands, crystallization of ice inside the bulk of the material could not be prevented at atmospheric pressure.

To the level of resolution and for type of printed scaffolds pursued in the experiments presented, no difference could be observed in the accuracy of imaging between using plunging in liquid ethane and in nitrogen. Both approaches allowed for faithful visualization of the alginate-based printed scaffolds. Nevertheless, it is documented that plunging in (undercooled) liquid ethane led in many scenarios to fewer artifacts in the highly hydrated samples, as it allowed for higher heat transfer [28, 29]. Only very high freezing velocities in combination with thin samples (up to 10 µm) led to the formation of vitreous ice at atmospheric pressure, whereas too slow heat transfer caused nucleation of ice and crystal growth [29]. Therefore, despite that the procedure with liquid nitrogen is overall faster, we recommend the use of the procedure with liquid ethane. The freezing velocities could especially have a clearly visible impact on imaging structures containing biological species (e.g. bacteria and cells) [28]. Note that the proposed use of a turbo-molecular pump and imaging without a conductive coating allows in general faster visualization and shorter investigation times than conventional cryo-SEM. As a result, the whole procedure of preparation by plunging in liquid ethane and SEM imaging of scaffold can be completed typically in less than 30 minutes.

In order to study the influence of solvent exchange on the structure of the printed scaffolds, we investigated plunge-freezing in liquid ethane after dehydration using ethanol/water series. After the first step of ethanol exchange using the 30% ethanol mixture, a slight shrinkage of the scaffold was observed based on light microscopic images (see Figure S4). This behavior can be explained by the formation of a denser network structure for calcium alginate gels in the presence of ethanol and a resulting volume contraction. For calcium-crosslinked alginate gels, the addition of ethanol to a solvent (water) up to a concentration of 24% was shown to influence the physical properties of the hydrogel, due to an alteration of the alginate chain (polymer coil) extension. Addition of alcohol at 15% or 24% led to the visible change from the homogeneous to a more heterogeneous and compact network structure [67]. After the subsequent dehydration steps and plunging into liquid ethane, however, no further change of the scaffolds dimension and 3D architecture was visible (see Figure S5). Importantly, no ice crystals were seen on top or in between the strands. Using pure ethanol for the last dehydration step in combination with drying by a soft tissue, the solvent was removed very efficiently. In the surface layer of the dehydrated samples, slight shrinkage was visible at higher magnification (Figure S3C). Inside the strands, a porous structure was observed (Figure S5C), with morphology and size range of the pores comparable to the porous structure found after plunge-freezing without dehydration step. This indicates a precipitation of ethanol crystals from the ethanol-polymer phase after solvent exchange and solidification by plunge-freezing. No differences to plunge-freezing of the original scaffolds in undercooled liquid ethane or liquid nitrogen were observed. Evidently, solvent exchange by ethanol and the solidification by plunge-freezing did not affect the precipitation and sublimation process of the shock-frozen calcium alginate gels in the studied scaffolds. Nevertheless, this preparation technique allowed to reduce the amount of the crystals on top and in between the strands, originating from the remaining water after tissue drying. However, due to the possibility of the more pronounced shrinkage during water/ethanol exchange and resulting morphological changes on the strands surface, this approach should be used with caution for the characterization of printed scaffolds.

### 3.3 Imaging of printed scaffolds with SEM

SEM imaging was performed at high and low vacuum conditions, depending on sample conductivity, in order to investigate the scaffolds without any conductive coating. No change in sample morphology was observed at 10 kV accelerating voltage during the imaging process both under high and under low vacuum. The maximal field of view was the same as in the cryo-SEM approach (10 mm by 10 mm), allowing to capture multiple printed strands and their crossing points in one image. To study the applicability of conventional SEM for faithful imaging of 3D printed scaffolds, the results are reported for the different preparation procedures:

#### 3.3.1 Air drying

Top-view and cross-section SEM images of the sample dried in air prior to imaging revealed a major structural rearrangement: large shrinkage of the printed strands both in x/y- and z-direction was observed in Figure 4B-D. The initial shape of the strands, visible on the LM images (Figure 4 A) was heavily altered, while only the overall shape of the native hydrated construct was maintained (Figure 4 B); the oval cross-section of the stands was greatly flattened (Figure 4 C and D). The strand surface appeared rough, with visible morphological changes and the formation of the artificial rims (see Figure S3D). In conclusion, the shape of the scaffolds changed radically and reliable information about the internal hydrogel structure cannot be deduced. While dehydration progressed, the condensation and densification of the hydrogel components occurred (see Figure 4D). The water loss led to deprivation of OH groups by the functional groups of alginate. The flexible polysaccharides chains became rigid due to the loss of the glycosidic bonds which provide conformational flexibility in the hydrated material. The viscosity of the hydrogel gradually increased, and eventually a solid state was reached [64].

**Figure 4.**
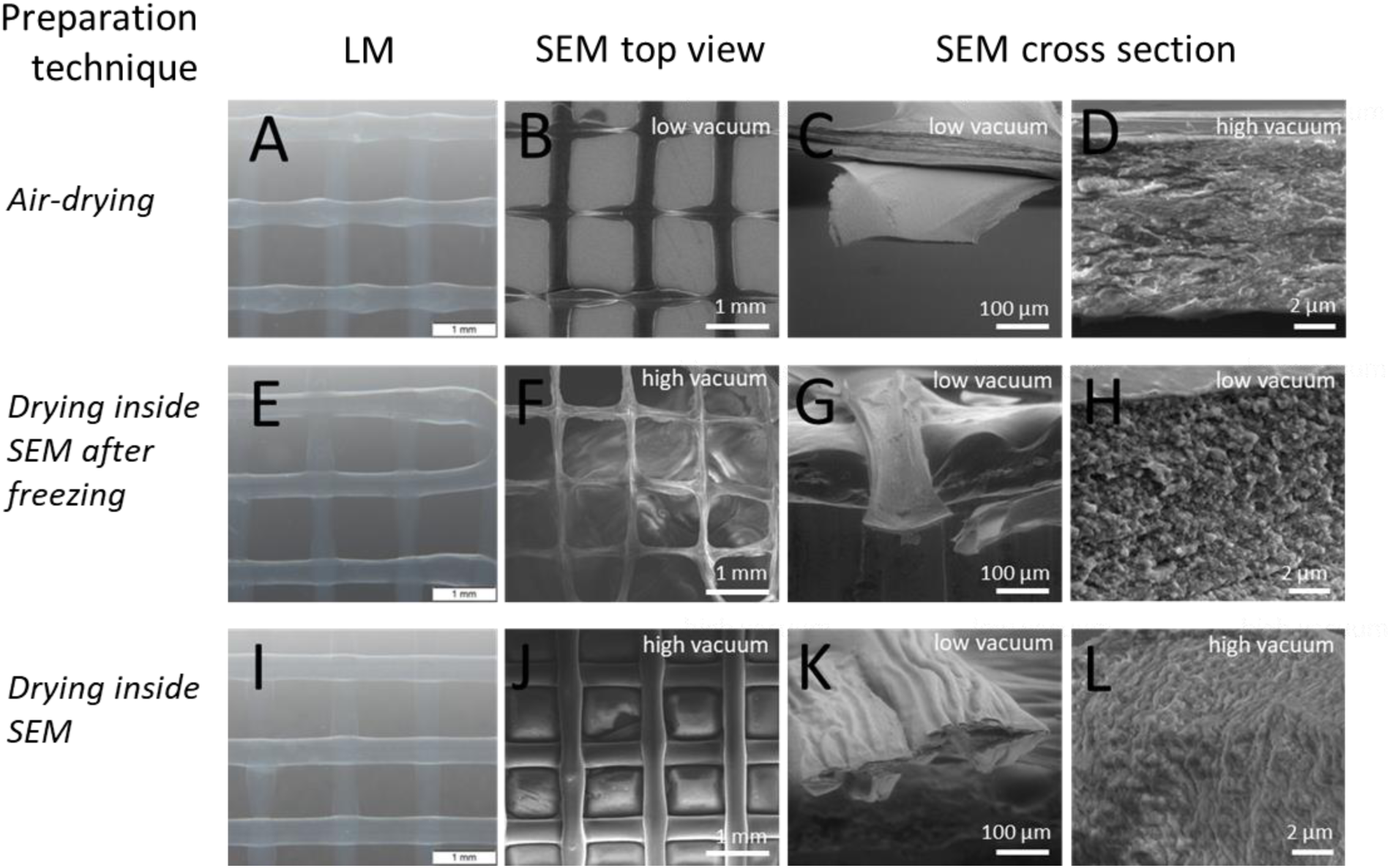
Conventional SEM investigation with different drying procedures: Light microscopic images of 3D printed scaffolds as received (A, E, I). SEM images after sample preparation by air-drying (B: top view; C-D: cross-section), drying inside SEM after storage in the freezer at T = 253 K/-20°C (F: top view; G-H: cross-section) and drying inside SEM (J: top view; K-L: cross-section). Drying leads to artifact formation in the form of a major reorganization of the structure, visible as shrinkage and densification/collapse.

#### 3.3.2 Drying onside the SEM after freezing

To analyze whether the artifacts caused by slow drying of hydrated scaffolds at ambient conditions can be avoided by sample freezing, scaffolds were initially frozen at T = 253 K/-20°C and subsequently dried inside SEM at T = 293 K/20°C under high vacuum conditions. These samples showed an extensive shrinkage of the printed strands (see Figure 4 F-H), even more pronounced than in the case of the air-dried scaffolds. The surface of the strands displayed morphological features resulting from the shrinkage (Figure S3E). These observations can be explained by the precipitation of ice during freezing and the resulting densification of the polymer phase during phase separation. We expect that the ice melted inside the SEM, which led to the slow evaporation of water between the polymer phase. These results indicate that the commonly used preparation technique of hydrogel printed scaffolds by freeze-drying is not recommended, as it leads to a major loss of the information about the real structure of the scaffold.

#### 3.3.3 Drying inside the SEM of a native sample

Interestingly, drying inside the SEM without preceding freezing led to more accurate preservation of the 3D structure (see Figure 4J-L). The top view images display a well-defined architecture, with no visible alteration in the strand size when compared to light microscopy images. We suppose that stabilization of the scaffolds structure in x- and y-direction can be explained by a high speed of water evaporation during drying inside the SEM chamber. In contrast to the frozen scaffolds, which are quite stiff after precipitation of ice, the as-prepared scaffolds stick well to the substrate and dry homogeneously during continuous evaporation of water inside SEM. However, in the z-direction, the loss of the original strand structure and the collapse of round-shaped geometry were observed (see Figure 4K). Parallel rims were visible on the strand surface at higher magnifications, clearly indicating strands shrinkage (Figure S3F).

### 3.4 Imaging of printed scaffolds in wet mode (ESEM)

Next, scaffolds were imaged using the environmental SEM, as this technique does not require any sample preparation steps. Due to the limited field of view by using a pressure-limiting aperture (diameter 500 µm), only a small area of the scaffolds can be visualized at once (max. 450 um × 450 µm). Therefore, the stitching of several images is necessary to characterize the 3D architecture of printed scaffolds (see Figure 5 A-B), which makes the approach time-consuming.

**Figure 5.**
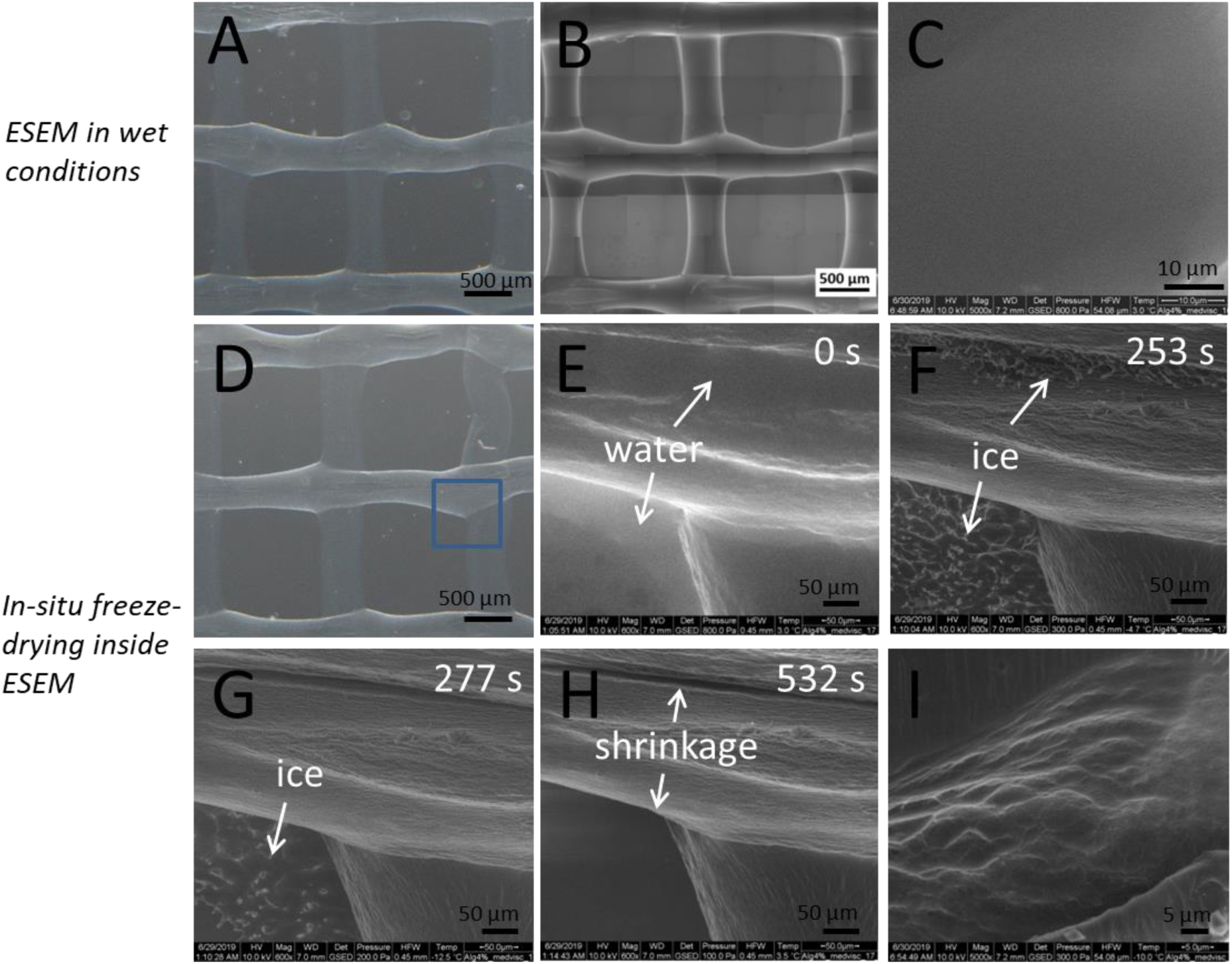
Environmental SEM investigation: Light microscopic image of 3D printed scaffolds as received (A, D). ESEM in wet conditions of scaffold in native hydrated state (B: top view after stitching of 70 ESEM images; C: cross-section) and in-situ freeze-drying inside ESEM (E-H: top view of the sample area marked by a blue square on 4D, E: in initial wet state showing water on top and next to the strands, F: after solidification and formation of ice crystals on top and next to the strands, G: after ice sublimation on top of the strand, H: after thawing and shrinkage; I: cross-section showing a porous structure after in-situ freeze-drying inside SEM at T = 263K/-10°C). ESEM in the wet state provides a well-preserved structure of the native hydrated construct.

Samples directly after washing were investigated in the water vapor equilibrium at T = 276 K/3°C and p = 800 Pa. In these conditions, no sample dehydration was expected. As shown in Figure 5B, the overall architecture of the scaffolds was well preserved with no signs of shrinkage. A smooth surface of the strands, without extensive wrinkles that would indicate drying, was captured (Figure 5E and Figure S3G). In comparison with other approaches, the surface morphology was by far best preserved. Cross-sections of the scaffolds in wet conditions were prepared with a blade. As shown in Figure 5C, a dense hydrogel structure, without any artificial porosity is visible. Note that special care was required for proper cross-sectional sample preparation as cutting in a wet state can easily alter the internal structure of the soft material. Apart from the limited field of view, ESEM had a clear advantage in providing exceptionally accurate representations of the printed structures.

The use of ESEM opened the possibility to perform an experiment of in-situ freeze-drying. The sample was cooled while simultaneously lowering the water vapor pressure inside the SEM chamber, according to the pressure/temperature phase diagram of water, by following the line of equilibrium between the liquid/solid and gaseous phase. The images, acquired in a period of 8 minutes and 52 seconds are shown in Figure 5 E-H. In this way, the morphological alterations of the samples following temperature and pressure changes were analyzed and structural changes that occurred during freeze-drying were continuously observed. Starting at T = 276 K/3°C and p = 800 Pa (Figure 5E), both temperature and pressure were decreased until the hydrogel scaffold solidified at T = 268 K/-5°C and p = 300 Pa (Figure 5F). No shrinkage of the strands was observed. Excess water crystallized on top and next to the strands (Figure 5F). Immediately after solidification, the ice from the strand surface sublimated in the vacuum (Figure 5G). Thawing of the samples to T = 276 K/3°C and p = 100 Pa leads to a shrinkage of the strands (see Figure 5H and Figure S3H). A cross-section image of a strand after in-situ freeze-drying at T =263 K/3°C and p = 300 Pa shows the formation of a porous structure in the strands due to ice precipitation and sublimation (see Figure 5I). The obtained results confirmed that freeze-drying of the samples before or during SEM imaging should be avoided in order to minimize artifacts and obtained faithful visualization of the hydrogel-based printed material.

### 3.5 Comparison of the tested approaches

We have performed imaging of 3D printed alginate-based samples in different microscopy modes (cyro-SEM, SEM and ESEM) after applying several preparation techniques to visualize the impact of sample handling on the accuracy of imaging. Our aim was to spot the possible inaccuracies and artifacts connected with the different approaches and to recommend the most suitable procedure for faithful visualization of printed hydrogel-based scaffolds. The main features associated with the distinct procedures presented in this study are summarized in Table 1.

**Table 1.**
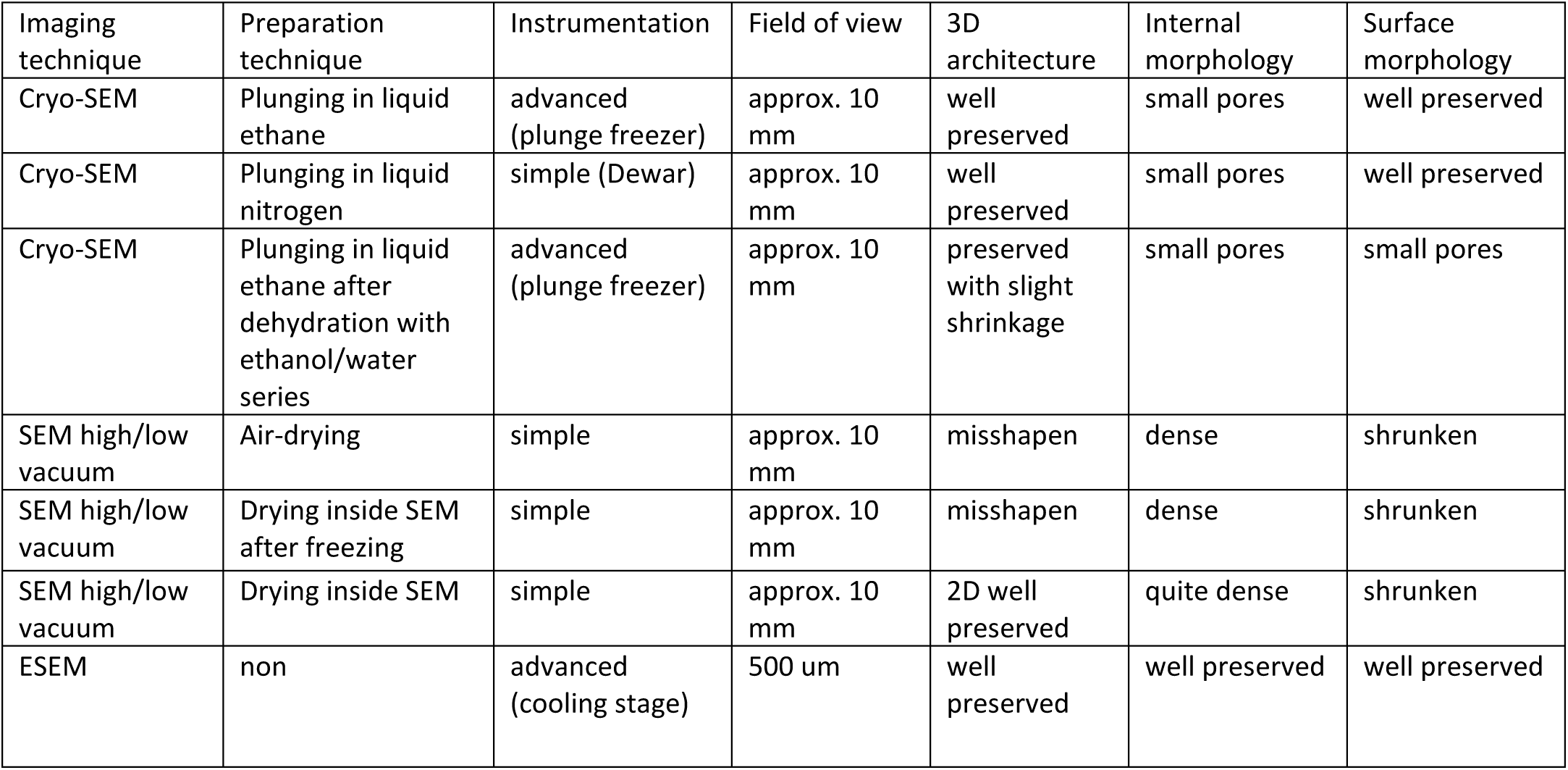
Comparison of different imaging modes and preparation techniques for SEM characterization of 3D printed hydrogels

| Imaging technique | Preparation technique | Instrumentation | Field of view | 3D architecture | Internal morphology | Surface morphology |
| --- | --- | --- | --- | --- | --- | --- |
| Cryo-SEM | Plunging in liquid ethane | advanced (plunge freezer) | approx. 10 mm | well preserved | small pores | well preserved |
| Cryo-SEM | Plunging in liquid nitrogen | simple (Dewar) | approx. 10 mm | well preserved | small pores | well preserved |
| Cryo-SEM | Plunging in liquid ethane after dehydration with ethanol/water series | advanced (plunge freezer) | approx. 10 mm | preserved with slight shrinkage | small pores | small pores |
| SEM high/low vacuum | Air-drying | simple | approx. 10 mm | misshapen | dense | shrunk |
| SEM high/low vacuum | Drying inside SEM after freezing | simple | approx. 10 mm | misshapen | dense | shrunk |
| SEM high/low vacuum | Drying inside SEM | simple | approx. 10 mm | 2D well preserved | quite dense | shrunk |
| ESEM | non | advanced (cooling stage) | 500 μm | well preserved | well preserved | well preserved |

Depending on the approach, we observed different levels of the preservation of the overall shape of the scaffolds, material structure and strands surface morphology. Cryo-SEM of the samples after plunging in undercooled liquid ethane or liquid nitrogen did not lead to substantial changes and resulted in only slight changes in the internal strand morphology and surface structure. Dehydration of the hydrogels by using a series of ethanol with increasing concentration led to a slight shrinkage during the first step of dehydration, giving no benefit to plunging the hydrogels scaffold in their native state. Drying inside the SEM resulted in a well preserved 2D structure, but in a strong shrinkage in z-direction due to the densification of the hydrogel by evaporation of water. Air-drying and drying inside SEM after freezing led to strong shrinkage in all dimensions and poor preservation of the 3D architecture. Drying and freeze-drying techniques heavily altered structural appearance, leading to inaccurate spatial information [64]. ESEM allowed visualization in the natural hydrated state, while the true hydrogel organization was fully maintained. However, it is limited by a small field of view (500 um × 500 um); as a consequence, the stitching of multiple images to visualize a bigger area of interest is required, which reduces the practicality of this technique. Therefore, we suggest a plunge-freezing procedure in liquid ethane, or alternatively in liquid nitrogen, as an optimal preparation technique to preserve the 3D architecture and morphology, with minimized artifacts, of the printed hydrogel-based scaffolds.

Using the recommended approach and a home-made setup (see Methods section), we performed further studies to test the applicability of the SEM imaging for 3D printing studies. Scaffolds with different ink compositions (see Table S1) were printed, visualized and characterized. Accurate information about size, shape, surface texture and internal structure of the printed constructs could be successfully obtained. For a more detailed analysis and discussion, we refer the reader to the Supplementary Information.

## 4 Conclusions

High-resolution SEM imaging is an excellent tool to advance the investigation of macro- and microscale features of printed constructs, greatly contributing to the developing field of biofabrication. However, the preservation of the native morphology of the printed material while imaging is not trivial. This study compared different approaches to prepare hydrogel-based printed samples for SEM imaging. Based on the performed comparison, the 3D architecture and morphology of printed hydrogels was best preserved by ESEM imaging of native scaffolds and by cryo-SEM after plunging in liquid (undercooled) ethane or nitrogen. Taking into consideration time efficiency and accessible field of view, we recommend cryo-SEM after plunging in liquid ethane (or nitrogen) for faithful visualization of hydrogel printed structures. We propose here a clear protocol of a fast and straightforward sample preparation for SEM imaging. It leads to high-quality cryo-SEM imaging, when used in combination with a home-made cryo-SEM sample holder and SEM equipped with a turbo-molecular pump. The results obtained for alginate-based scaffolds can be extended to a large range of hydrogels and the proposed protocol can be easily adapted to other types of printed materials.

We suggest that cryo-SEM, applied with special caution regarding sample preparation, should be included in the ink development studies as a powerful complementary tool to light microscopy. When used properly, it provides an accurate and detailed visualization of 3D architecture and surface morphology of printed scaffolds and single filaments, at a level difficult to obtain with other microscopic techniques.

## 5 Acknowledgment

The authors would like to thank Christine Hartmann for proof-reading the manuscript, Prof. Eduard Arzt for fruitful discussions.

This research did not receive any specific grant from funding agencies in the public, commercial, or not-for-profit sectors.

