## Supplementary Information for "Faithful Scanning Electron Microscopic (SEM) visualization of 3D printed alginate-based scaffolds"

1. Scaffold preparation

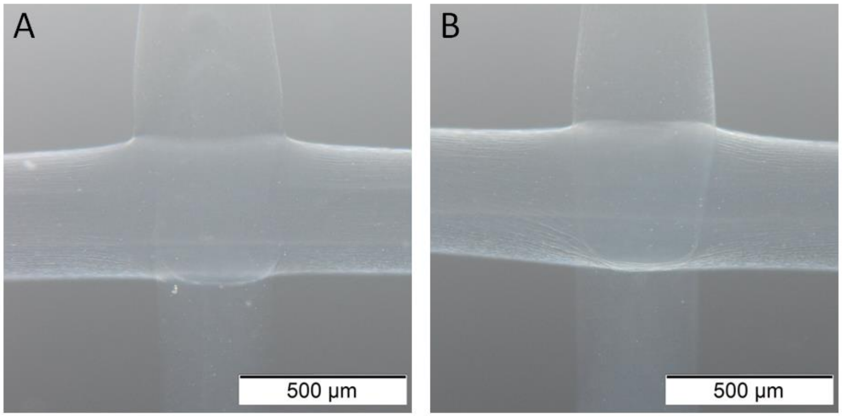


Figure S1. Scaffold structure observed after storage in the printing bath solution for 5 hours (A) and 11 days (B) prior to imaging. No substantial changes can be observed.

2. Results: Imaging based on different SEM modes

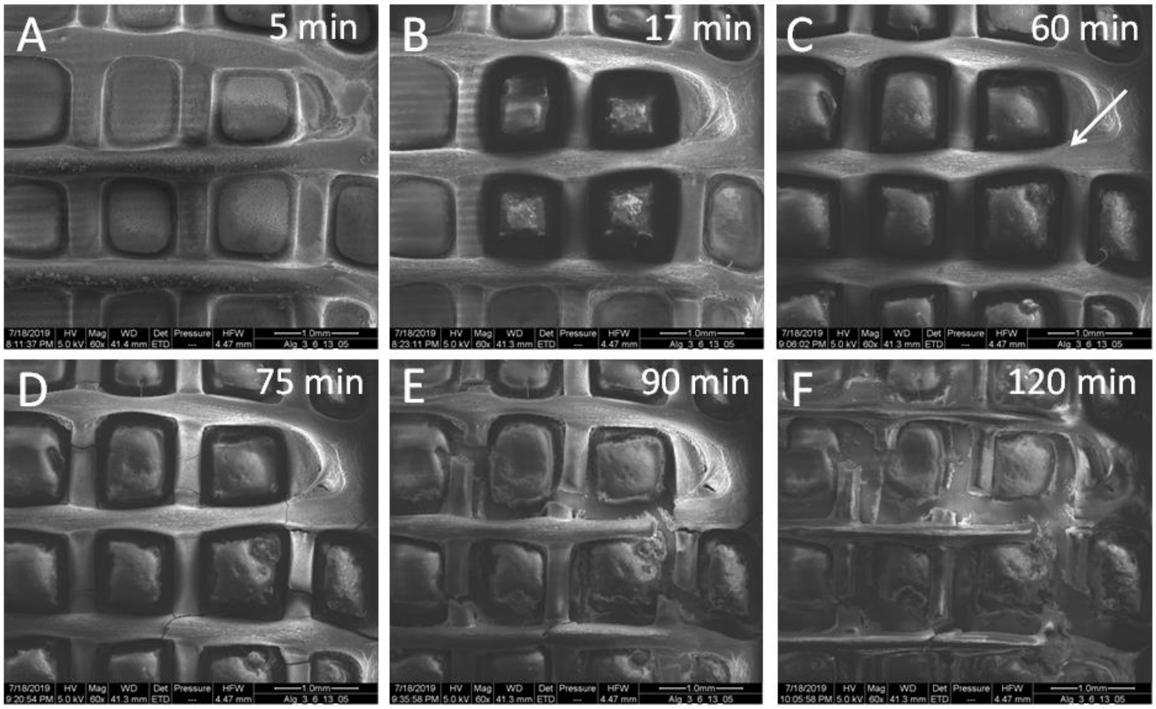


Figure S2. Cryo-SEM images of a printed sample (with additional post-printing crosslinking in 100mM $CaCl_2$) after plunge-freezing in liquid ethane and during thawing under high vacuum conditions. Time of imaging after plunging is indicated in the right bottom corner of each image. Crack formation starts approximately 60 minutes after vacuum initialization (see arrow in C).

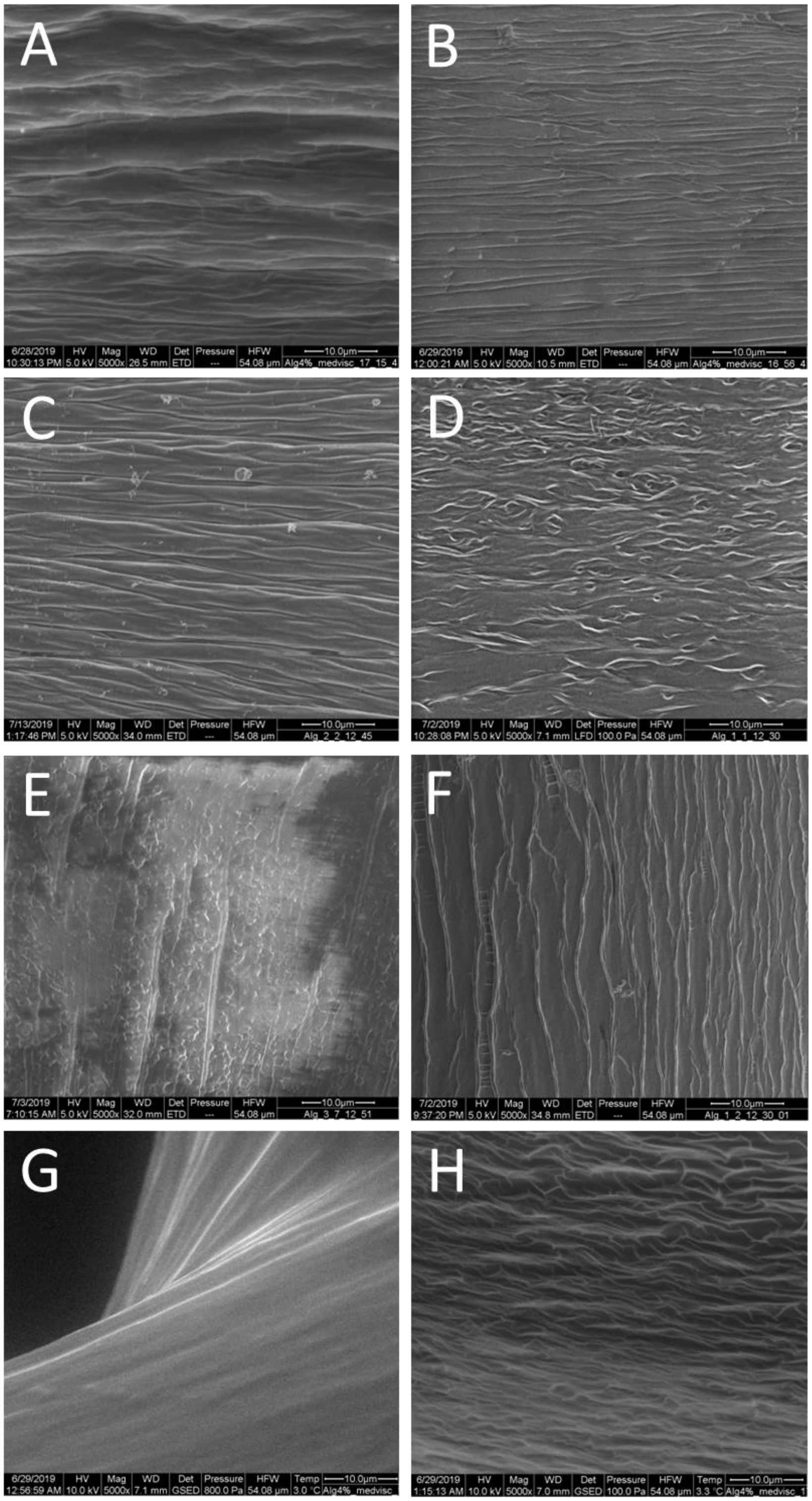


Figure S3. Comparison of the surface morphology for different preparation techniques: Plunge-freezing in liquid ethane (A), plunge-freezing in liquid nitrogen (B), plunge-freezing in liquid ethane after dehydration (C), air-drying (D), drying inside SEM after storage in the fridge at T = 253 K/-20°C (E), drying inside SEM (F), imaging in wet mode/ESEM (G) and in-situ freeze-drying inside ESEM (H).

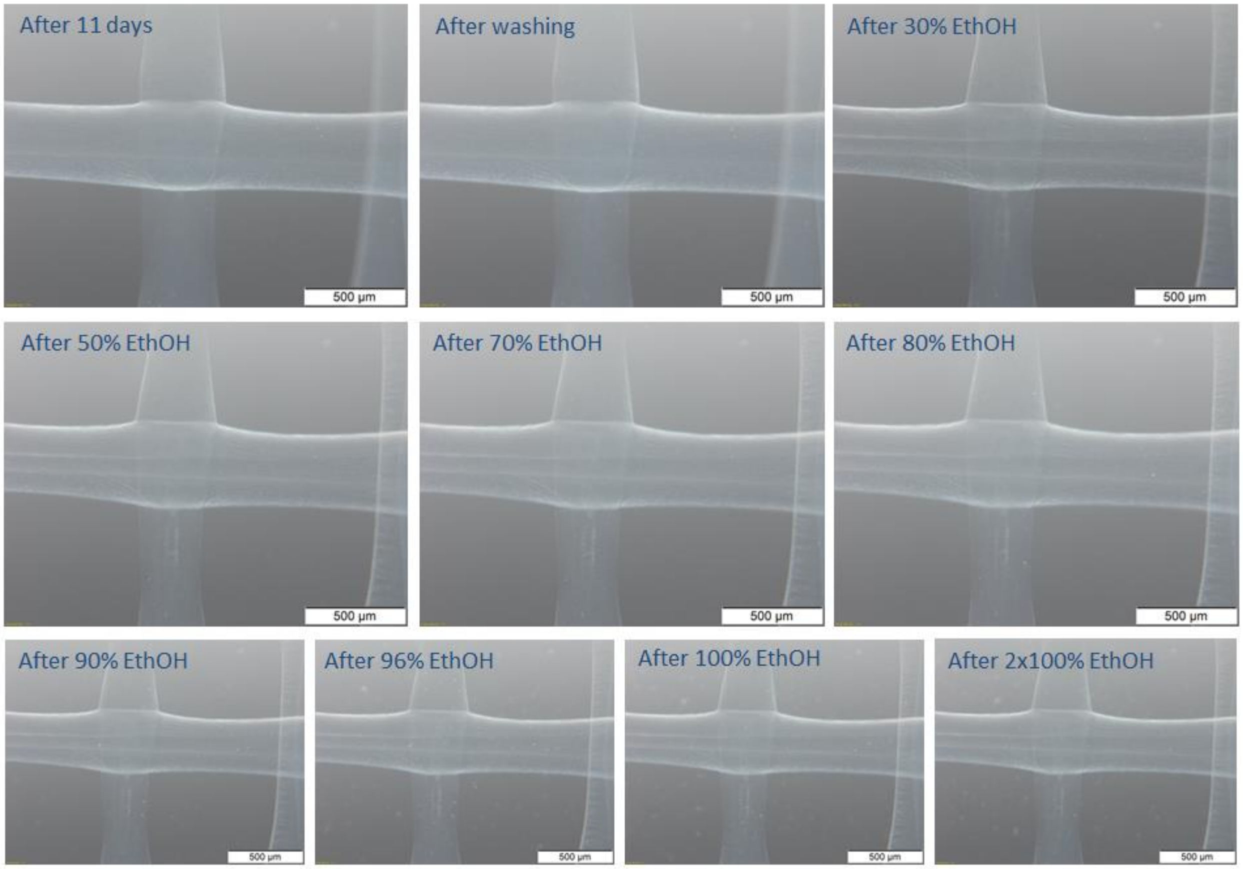


Figure S4. Light microscopy of the printed scaffold (stored for 11 days in the printing bath solution) after washing with demineralized water and dehydration using a series of ethanol/water mixtures with increasing ethanol concentration: Slight shrinkage of the strands can be observed after the first dehydration step (30% ethanol).

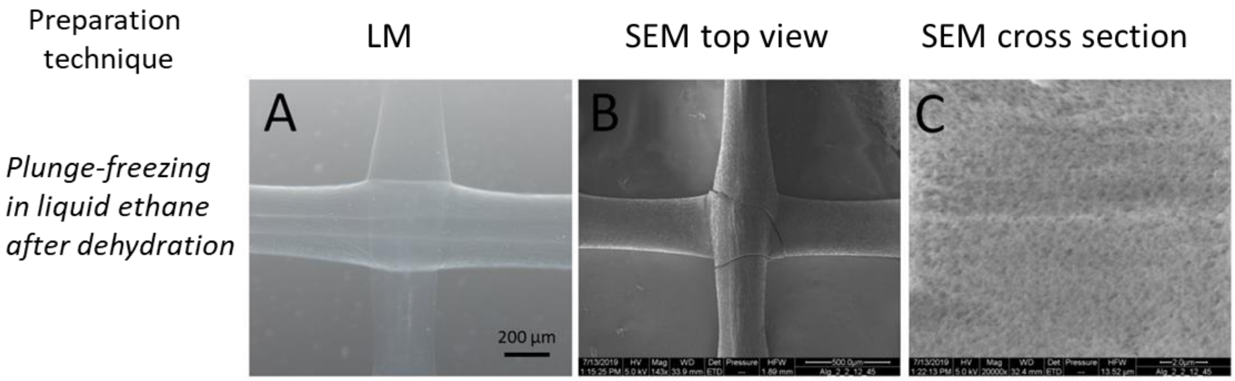


Figure S5. ESEM investigation: Light microscopic image of 3D printed scaffolds as received (A). ESEM images after sample preparation by dehydration with ethanol/water series followed by plunge-freezing in liquid ethane (B: top view; C: cross-section). The structure is not well-preserved, with the evidence of scaffold shrinkage, crack formation and artificial porosity inside the strand.

### 3. The applicability of the faithful SEM imaging for 3D printing field

#### 3.1 Ink preparation and scaffolds printing

Inks composed of 4% (w/v) solution of alginic acid sodium salt (medium or high viscosity) in Milli-Q water were prepared. The samples were physically mixed with a spatula, followed by the use of NeoLab multifunctional mixer (few hours or overnight) until homogeneous solutions were obtained. Solutions were loaded into 3 ml or 10 ml printing cartridges, sonicated at T = 323 K/50°C (30 – 45 minutes) and centrifuged at 2000 rpms (at T = 298 K/25°C for 25 minutes; only the inks containing high viscosity alginate) to remove air bubbles that have appeared while mixing.

Bath solutions were prepared by mixing in 1:1 volume ratio: 0.05% (w/v) chitosan in 2% (v/v) acetic acid solution with 10 mM or 100 mM $CaCl_2$, leading to the final printing bath compositions described in Table 1. Additional, post-printing treatment with 100 mM $CaCl_2$ was introduced for improved stability of some of the scaffolds.

In total, 2 different ink formulations, 2 bath solutions, and 3 crosslinking conditions were used in this study, leading to 4 different scaffold types (see Table S1).

Table S1: Abbreviations of scaffold types used in the study, with indicated ink compositions, bath solutions and crosslinking conditions.

| Scaffold type abbreviation | Ink formulation | Final bath solution composition | Post-printing crosslinking |
|---|---|---|---|
| AlgM | Medium viscosity alginate 4% (w/v) | Chitosan 0.05% (w/v) and 5 mM $CaCl_2$ | Not applied |
| AlgM/100 $CaCl_2$ | Medium viscosity alginate 4% (w/v) | Chitosan 0.05% (w/v) and 5 mM $CaCl_2$ | 100 mM $CaCl_2$ |
| AlgH5 | High viscosity alginate 4% (w/v) | Chitosan 0.05% (w/v) and 5 mM $CaCl_2$ | Not applied |
| AlgH50 | High viscosity alginate 4% (w/v) | Chitosan 0.05% (w/v) and 50 mM $CaCl_2$ | Not applied |

Alginate-based 2-layers grid-like constructs were printed using square design with ca. 1 cm x 1cm dimensions (1.5 mm strand distance), and perpendicular layers placement. Medium viscosity alginate (AlgM) constructs were printed with 20 – 40 kPa pressure and 10 mm/s printing speed. High viscosity alginate (AlgH) constructs were printed with 80 kPa pressure and 10 mm/s printing speed. Some AlgM scaffolds were additionally crosslinked with 100 mM $CaCl_2$ for at least 2 hours to further increase stability (AlgM/100 $CaCl_2$). Printing was performed directly into the bath solution (see Table 1), on the glass slides fixed with type stripes in 6 well-plates or directly into the plate, using 250 µm inner diameter conical needle (gauge 25), with cartridge temperature T = 313 K/40°C and bath temperature T ~ 308

K/35°C. Z-offset of the first layer was ca. 150 µm for medium viscosity alginate and 0 for high viscosity alginate, layer height was set to 180 µm for medium viscosity alginate and 200 µm for high viscosity alginate, to ensure a good connection between glass and consecutive printed layers. Printers BioScaffolder 3.2 (GeSiM, Germany) and Bio X (Cellink, Sweden) were used. Prior to imaging, samples were stored in bath solution or 100 mM $CaCl_2$ at T = 277 K/4°C for up to 11 days and no visible change of scaffold structure could be observed (Figure S1).

### 3.2 Statistical analysis

Strands measurements are reported as the mean ± standard deviation. Statistical differences were analyzed with InStat3 software. Unpaired t-test, with a two-tailed P value, was performed for comparison of strand width based on SEM and LM images for each scaffold (intersections and bridging strands, separately) and one-way Analysis of Variance (ANOVA), followed by post-hoc Tukey-Kramer test. Multiple Comparisons Test was performed for comparison of strand sizes in different scaffolds (intersections and bridging strands) measured based on SEM or LM images, separately. Based on the robustness of the ANOVA test, the violation of the assumption of homogeneity of variance was ignored. Significant differences are considered for $p < 0.05$.

### 3.3 Cryo-SEM after plunging in liquid ethane: detailed analysis

4 types of scaffolds were printed at the conditions described in Table S1. LM images of the scaffolds, presented in Figure S6, provide information about the general shape of the scaffold but do not allow to observe any details regarding surface morphology/topography of the scaffolds or single strands. SEM visualization (Figures S7, S14-S18) provides more detailed information.

#### 3.3.1. Imaging at lower magnification

We have used lower magnification imaging (original mag. 40x) to obtain information about printing quality and to perform measurements of the width of printed strands (Figure S7).

Figure S7 (low magnification) provides information about the quality of the ink: the extent of the collapsing of the layers is well visible. For AlgM sample (Figure S7E), without post-printing crosslinking, clearly overhanging structures indicate that the viscosity of the ink is too low to maintain the shape for a longer time. After crosslinking with 100 mM $CaCl_2$, the shape is stabilized, resulting in less overhanging (see Figure S7F). For samples printed with high viscosity alginate (AlgH5 and AlgH50, Figure S7G and S7H, respectively) the collapsing of material strands can be observed to a lesser extent. The viscosity is high enough for obtaining self-holding structures.

For AlgM and AlgM/100 $CaCl_2$ samples, the higher smoothness of the strands in comparison to AlgH5 and AlgH50 can be observed (Figure S7). For the samples with high viscosity alginate, the lack of material homogeneity is clearly visible as strands surface roughness. This inhomogeneity could be caused by improper mixing of the printed solution (due to the ink's high viscosity) or by not fully dissolved alginate. For AlgH50 sample, a flattening of the strands is found, which can be caused by too low z-offset or by hydrogel shrinking due to the incubation in the solution containing a higher concentration of calcium ions [1]. Additionally, the connection between strands of the consecutive layers is well-visualized for all the scaffolds.

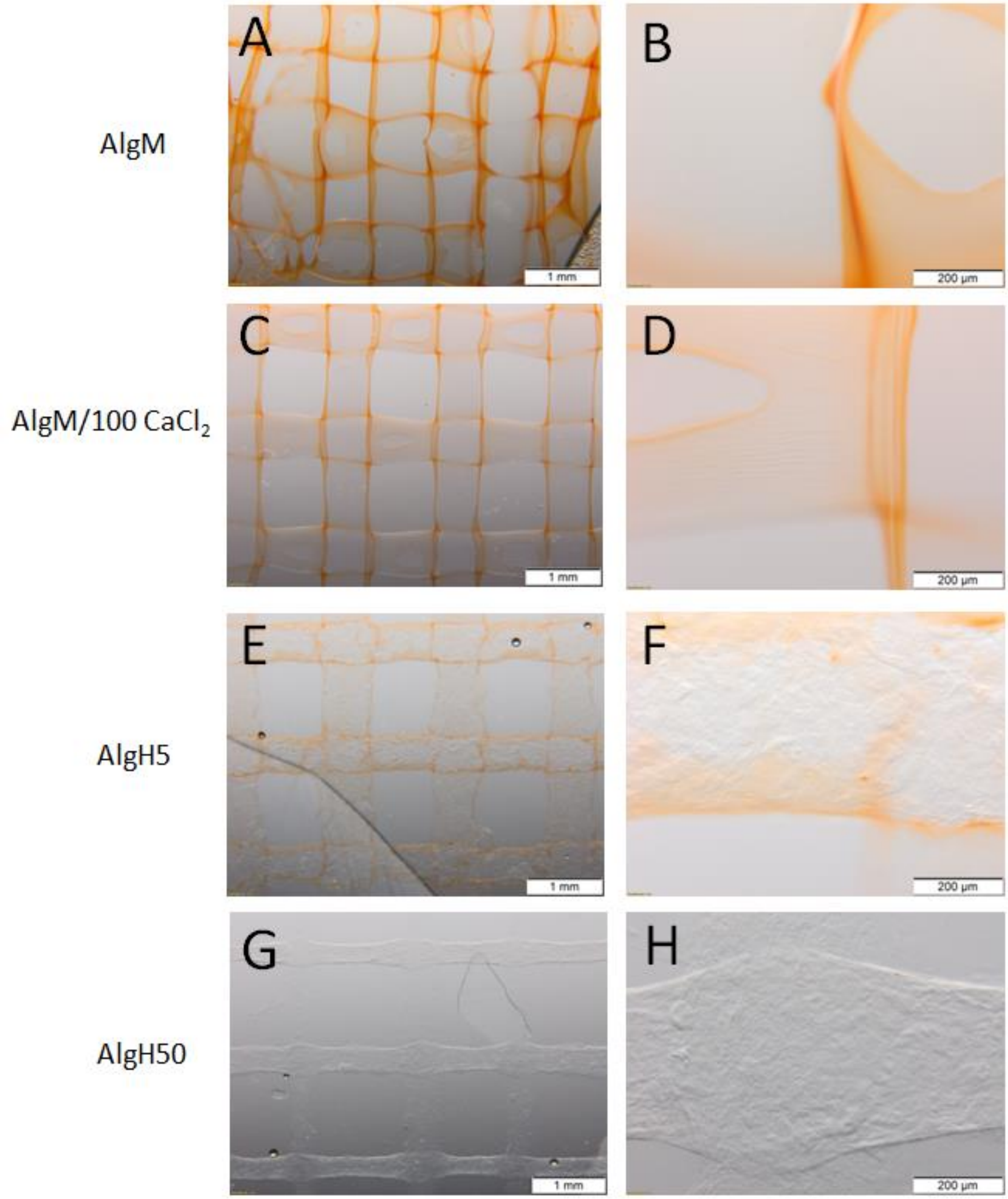


Figure S6. Light microscopic imaging of AlgM (A, B), AlgM/100 $CaCl_2$ (C, D), AlgH5 (E, F) and AlgH50 (G, H).

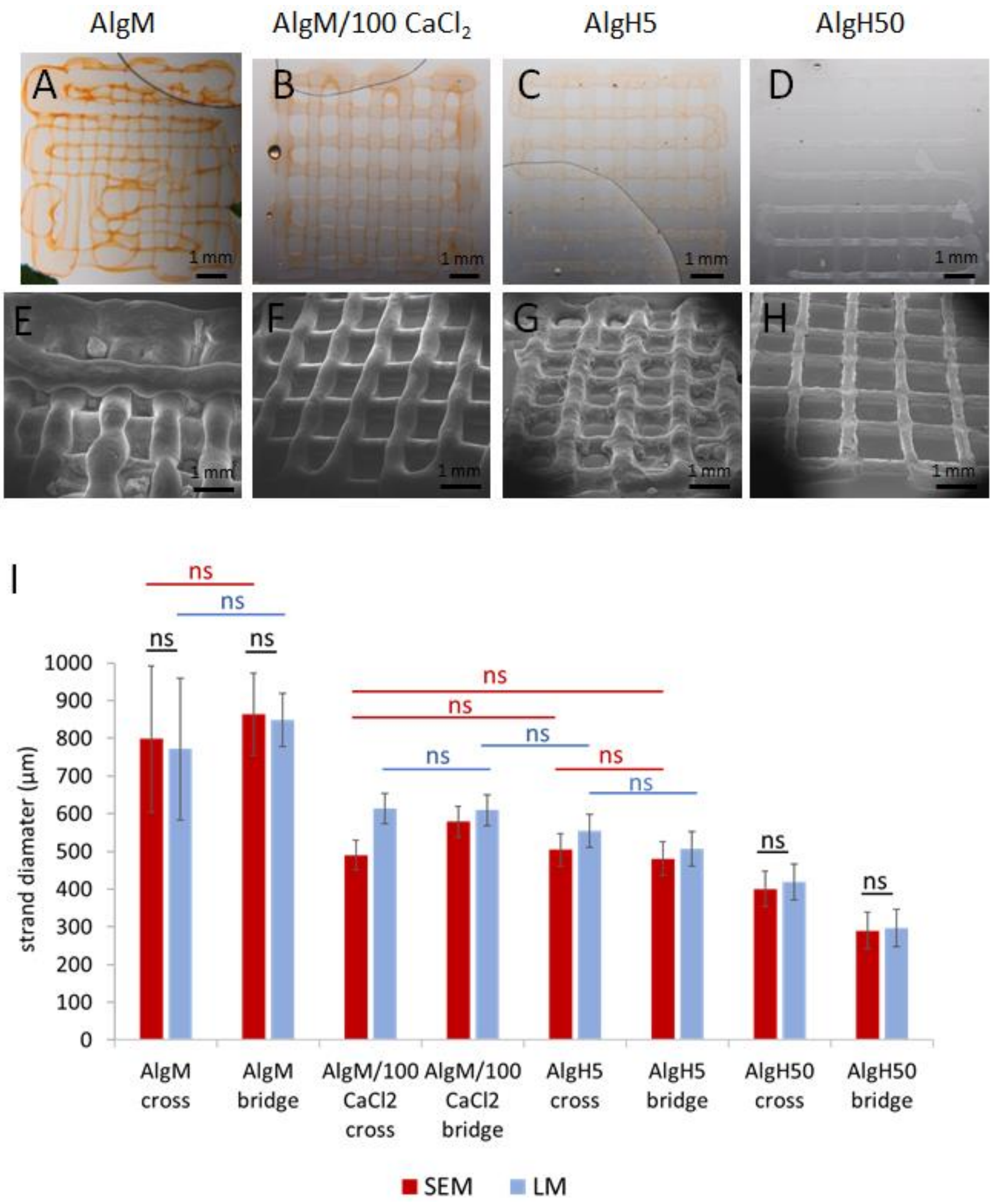


Figure S7. Light microscopic (A-D) and cryo-SEM images (E-H) at 40x original magnification of AlgM, AlgM/100$CaCl_2$, AlgH5 and AlgH50. Comparison of average diameter and standard deviation of the strand width measured by LM (in blue) and cryo-SEM (in red) for the crossings and bridges (I). Only not significant differences are marked on the figure (ns). All other differences were significant, with $p \leq 0.05$

The diameter of the strands was analyzed for top-view images acquired with LM and cryo-SEM at 40x magnification, in the part of the filament bridging two strands lying below (bridges) and in the crosslinking point of the strands from different layers (crossings) (See Figure S7 and Figures S8-S13 for exact measurements). For AlgM and AlgH50, the strand width values obtained using LM and SEM are comparable, no significant differences were detected. For the other two imaged scaffolds, strand measurement based on LM images gives higher values than based on SEM imaging. We hypothesize that this is connected to less exact visualization of the strand borders in the LM imaging.

The widest strands, based on both imaging techniques, were obtained for AlgM (SEM: 864µm±109µm for bridging parts of the fibers and 798µm±193µm for crossings with lower layer; LM: 848µm±71µm for bridges, 771µm±188µm for crossings). This could be explained by the collapsing of the layers for AlgM, causing a relative widening of the filaments. Additionally, some swelling while storing in the solution prior to imaging could have occurred. For medium viscosity alginate with additional post-printing treatment with 100mM $CaCl_2$, strand sizes were significantly smaller (SEM: 580µm±30µm bridges and 491µm±39µm crossings; LM: 609µm±29µm bridges and 613µm±28µm crossings), which can be assigned to the positive influence of additional crosslinking on the shape maintenance. AlgH5 strand size was smaller than AlgM and AlgM/100$CaCl_2$ based on both imaging techniques (SEM: 481µm±32µm bridges and 504µm±57µm crossings; LM: 506µm±29µm bridges and 559µm±54µm crossings). However, the difference between bridging parts of the fibers for AlgM/100$CaCl_2$ and AlgH5, as measured based on SEM, was not significant. As the printing settings for all the scaffolds where the same, ideally, the strand size should be the same for all the scaffolds. The wider strands in the bridging parts can be explained by more pronounced collapsing of the filaments in scaffolds with lower viscosity of alginate. Finally, the smallest strand dimensions were observed for AlgH50 scaffolds printed with high viscosity alginate into the crosslinking bath with higher calcium content (SEM: 290µm±48µm bridges and 401µm±62µm crossings, LM: 297µm±49µm bridges and 419µm±56µm crossings). We speculate that the higher extent and speed of crosslinking, due to the higher $CaCl_2$ content, can explain the smallest strand size. Additionally, we have observed the significant differences between crossing and bridging parts of the filaments for AlgM/100$CaCl_2$ based on SEM images, but not on LM images, and based on both imaging techniques for AlgH50. For other scaffolds, no differences in size between two measure regions of fibers were detected. The bigger size of bridges observed based on SEM for AlgM/100$CaCl_2$ can be explained by well visible overhanging, collapsing of structures, which is not so clearly visible based on LM pictures. For AlgH50, the crossing points are wider due to the merging of consecutive layers and diffusion of the material at the crossings, whereas the fibers in the bridges do not overhang but shrink, leading to the narrower size. In conclusion, SEM imaging gives a more accurate base for the strand size estimation and comparison, especially if the differences between scaffolds and changes in morphology are subtle.

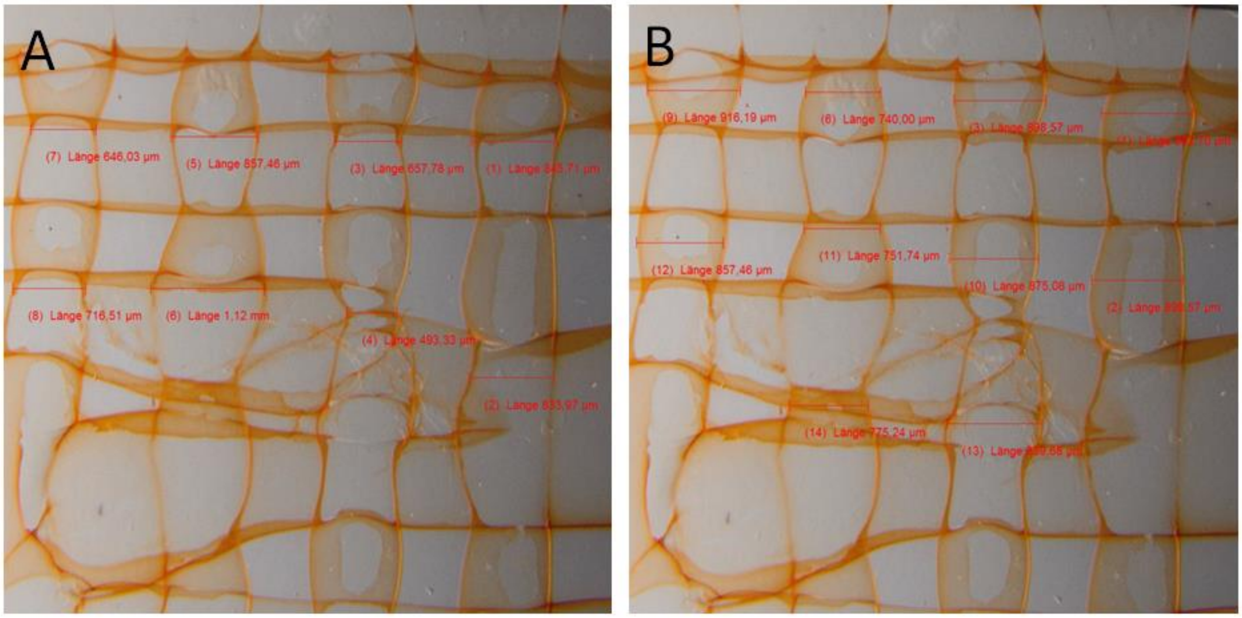


Figure S8. Strand size measurements based on light microscopic imaging of AlgM scaffolds (A: crossings, B: bridges).

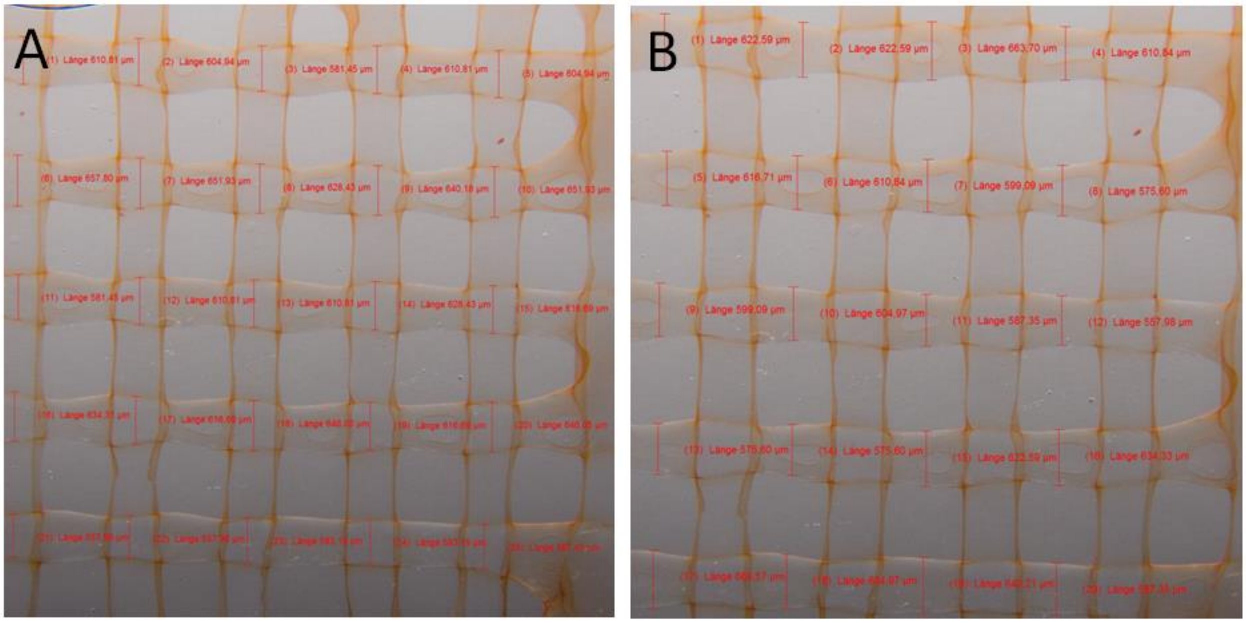


Figure S9. Strand size measurements based on light microscopic imaging of AlgM/100 $CaCl_2$ scaffolds (A: crossings, B: bridges).

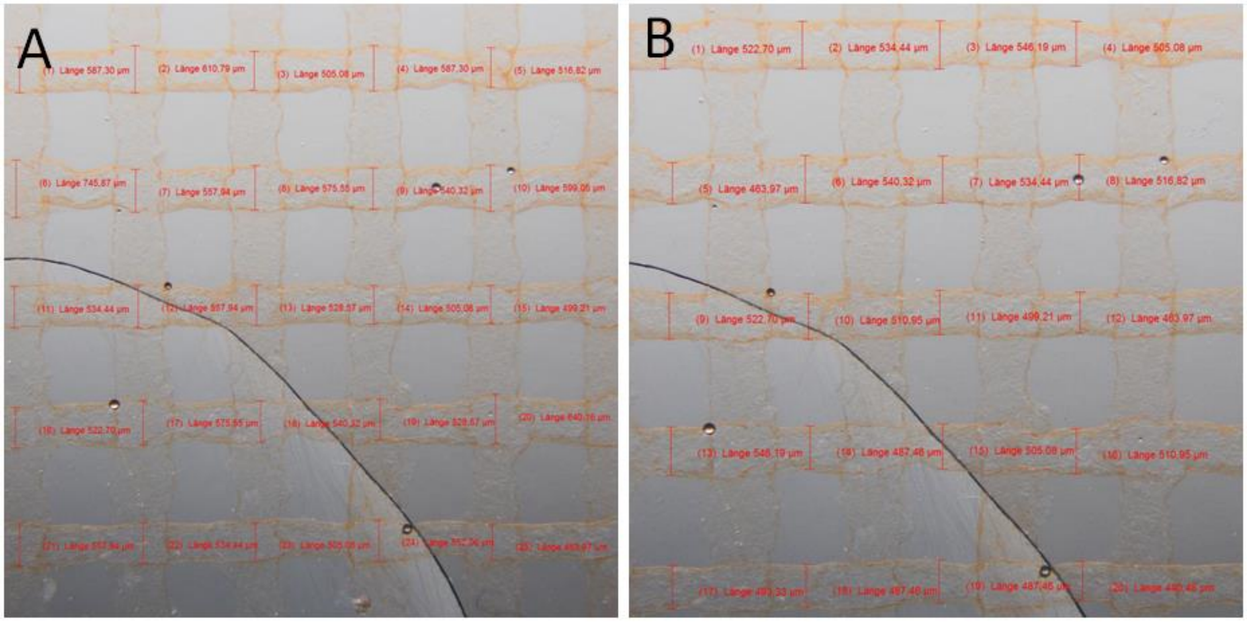


Figure S10. Strand size measurements based on light microscopic imaging of AlgH5 scaffolds (A: crossings, B: bridges).

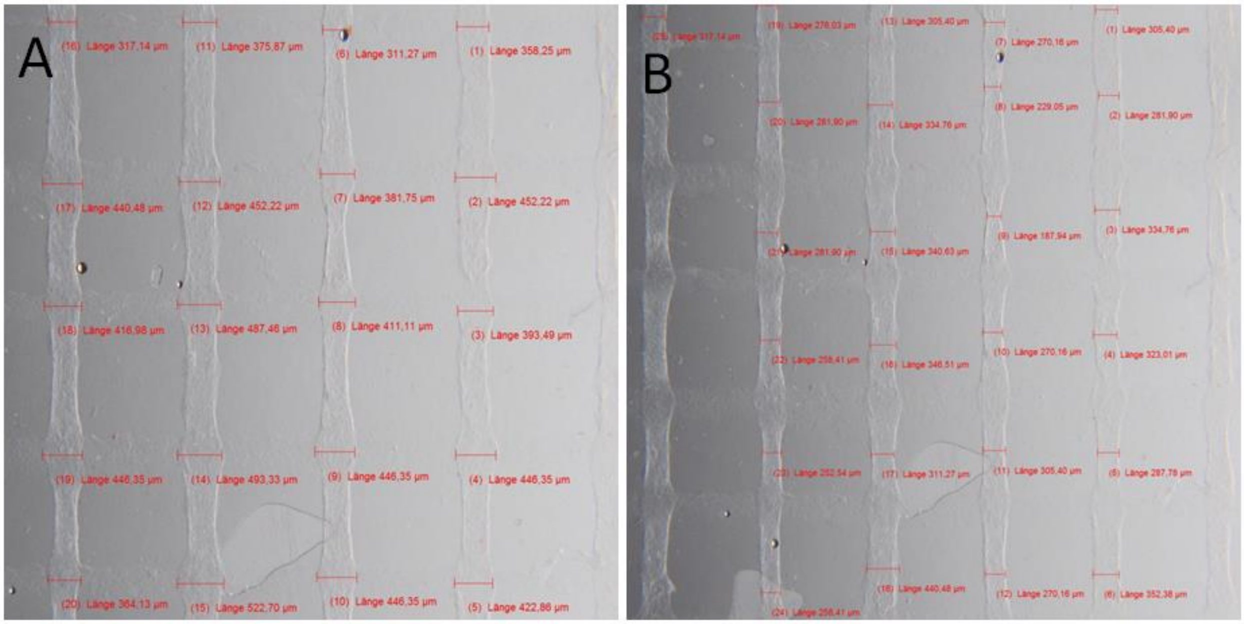


Figure S11. Strand size measurements based on light microscopic imagining of AlgH50 scaffolds (A: crossings, B: bridges).

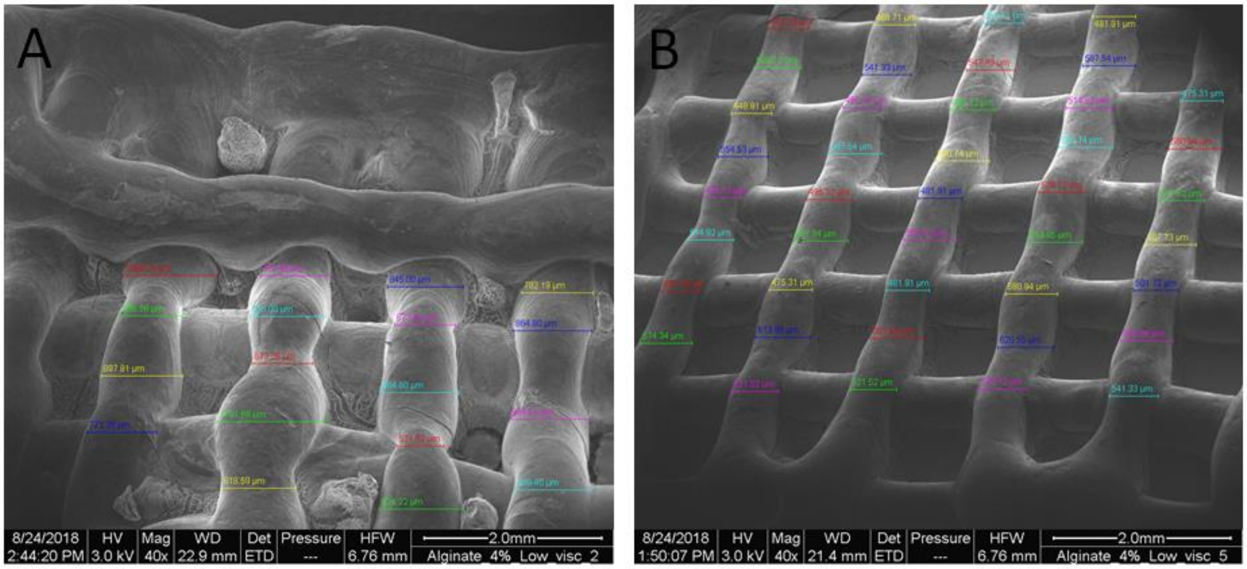


Figure S12. Strand size measurements based on SEM imaging of AlgM and AlgM/100 $CaCl_2$ scaffolds.

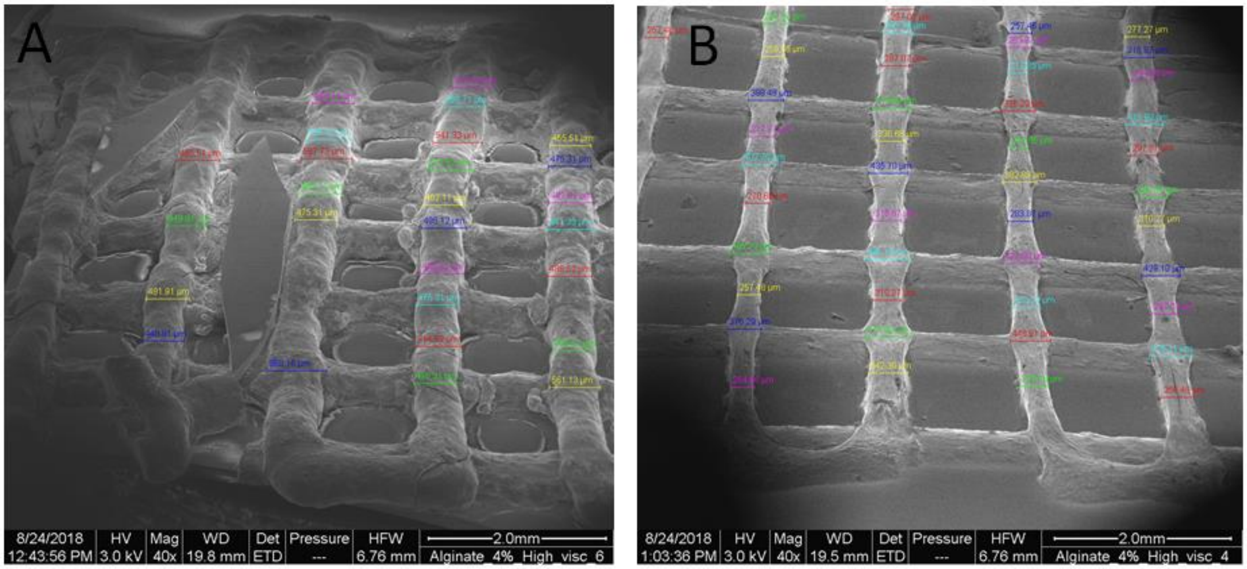


Figure S13. Strand size measurements based on SEM imaging of AlgH5 and AlgH50 scaffolds.

3.3.2. Imaging at higher magnifications

Cryo-SEM images at higher magnifications (original mag. between 100x and 5000x) were obtained with very good resolution (see Figure S14-S18). The strand shape, strand surface morphology, integration between consecutive layers, their fusion and the interaction between the bottom layer and printing surface are clearly visualized.

At low magnifications, strands printed with medium viscosity alginate (AlgM) reveal smooth surface, whereas ones printed with high viscosity alginate show clear inhomogeneity. This indicates better homogeneity of medium viscosity mixtures. However, at high magnifications (original mag = 5000x) a particle-like morphology is found for AlgM samples (Figure S14I and J). We assume that these particles are ice crystals that precipitate inside the hydrogel during the plunge-freezing process. In comparison, at AlgH strands surface, no ice crystals are visible, probably due to the higher viscosity of alginate and better bounding of water inside the network. Instead, small wrinkles on the surface can be noticed, indicating shrinkage (Figure S14 K, L). For AlgH50, the strands are partially cracked (Figure S14 D). Most probably, this effect is caused by the different thermal expansion of the scaffolds and the substrate (glass or plastic), or by the slow warming-up leading to ice sublimation while imaging under high vacuum.

For sample AlgM/100$CaCl_2$, a cross-section analysis by freeze-fracture under liquid nitrogen after mounting the sample onto the cryo-SEM holder was used to visualize the dimensions and internal structure of the printed fiber (Figure S14 M). The observed strands have a homogenous internal structure (no artificial porosity) and are nearly round shape. These indicate no flattening was caused by printing, neither by the preparation of the scaffolds by plunge-freezing.

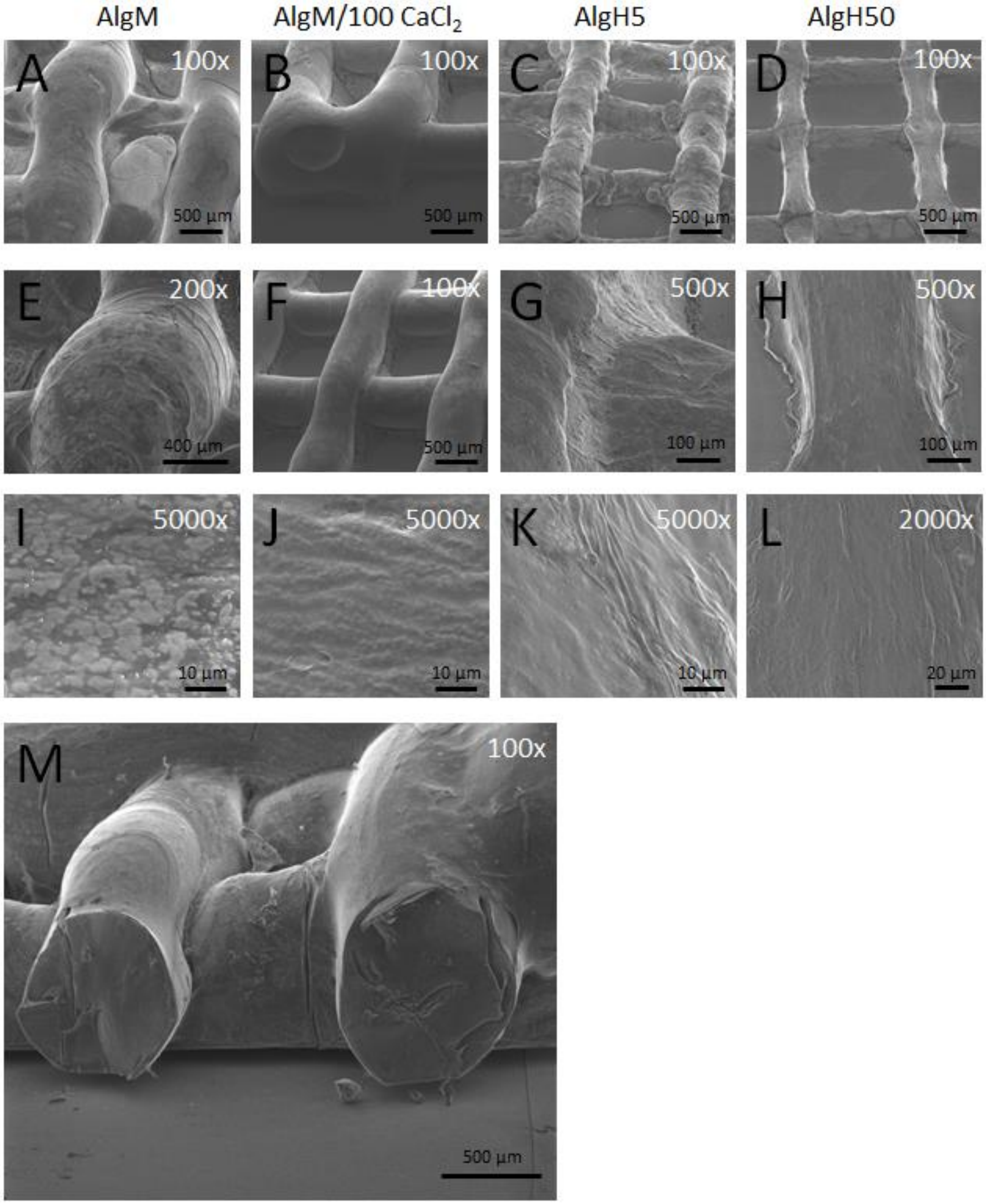


Figure S14. Cryo-SEM images at different magnifications (indicated in white) of AlgM (A, E, I), AlgM/100$CaCl_2$ (B, F, J) AlgH5 (C, G, K) and AlgH50 (D, H, L) and cross-section of AlgM/100$CaCl_2$ (M)

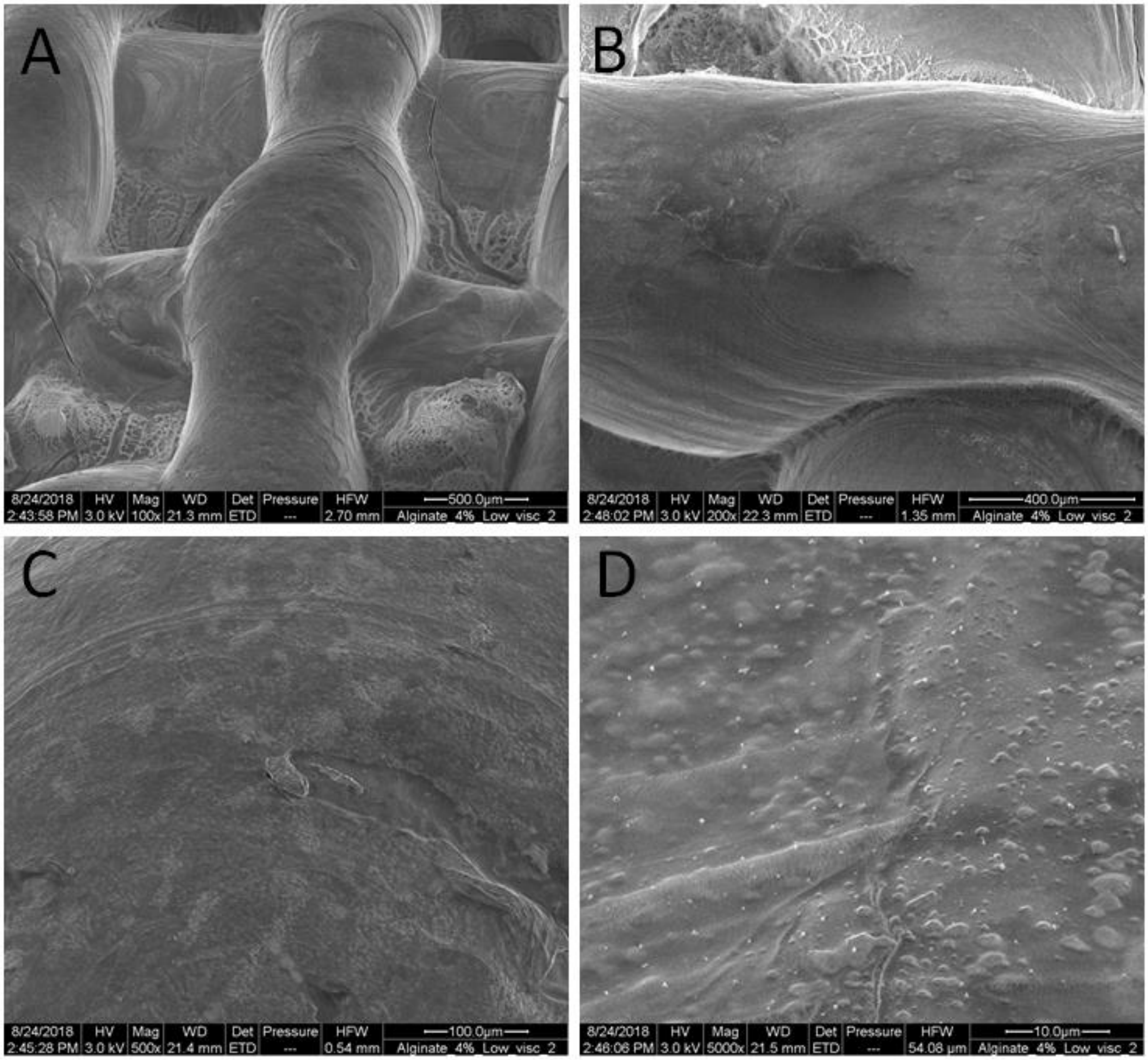


Figure S15. Morphological characterization of AlgM by SEM (A-D).

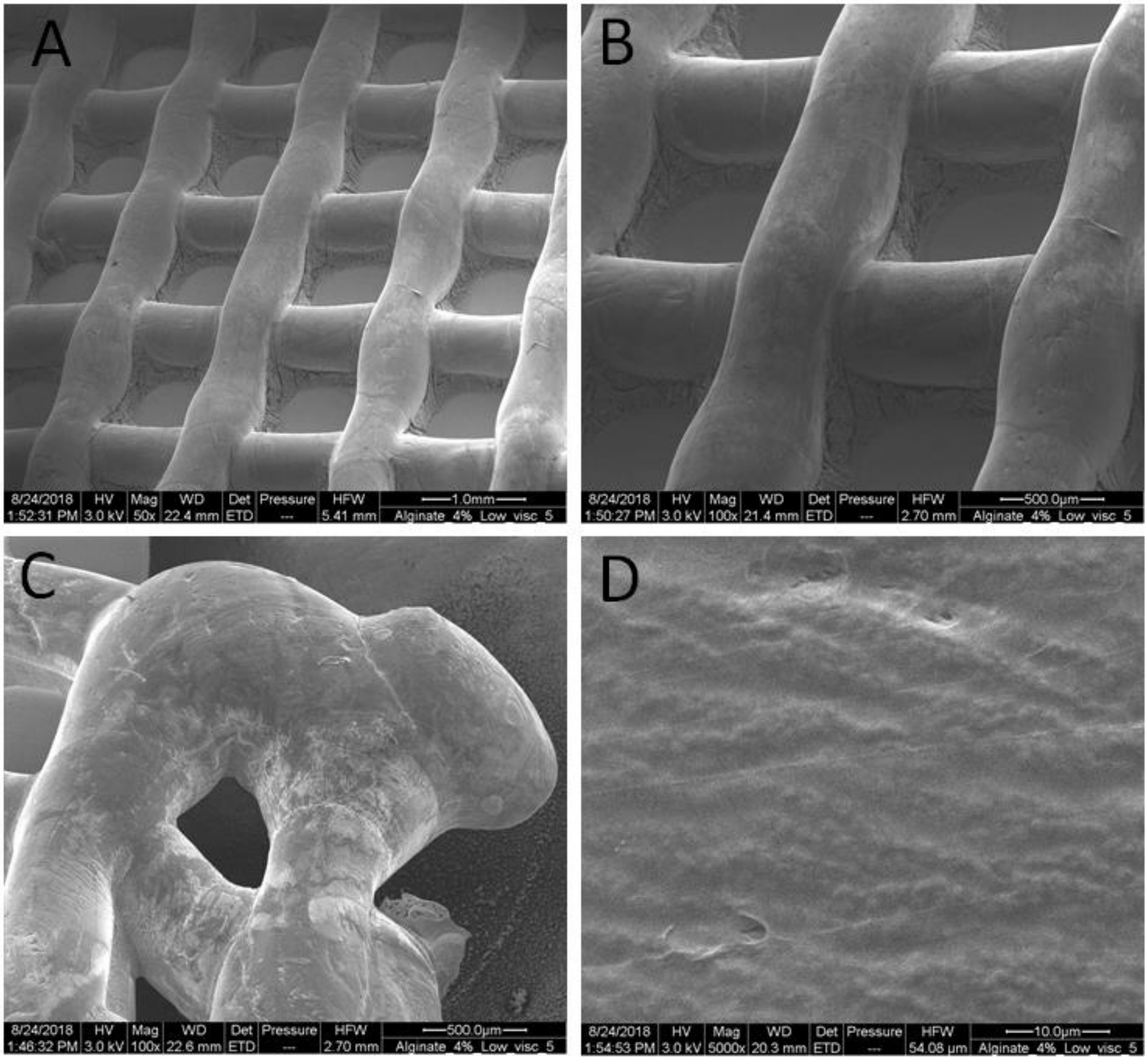


Figure S16. Morphological characterization of AlgM/100 $CaCl_2$ by SEM (A-D).

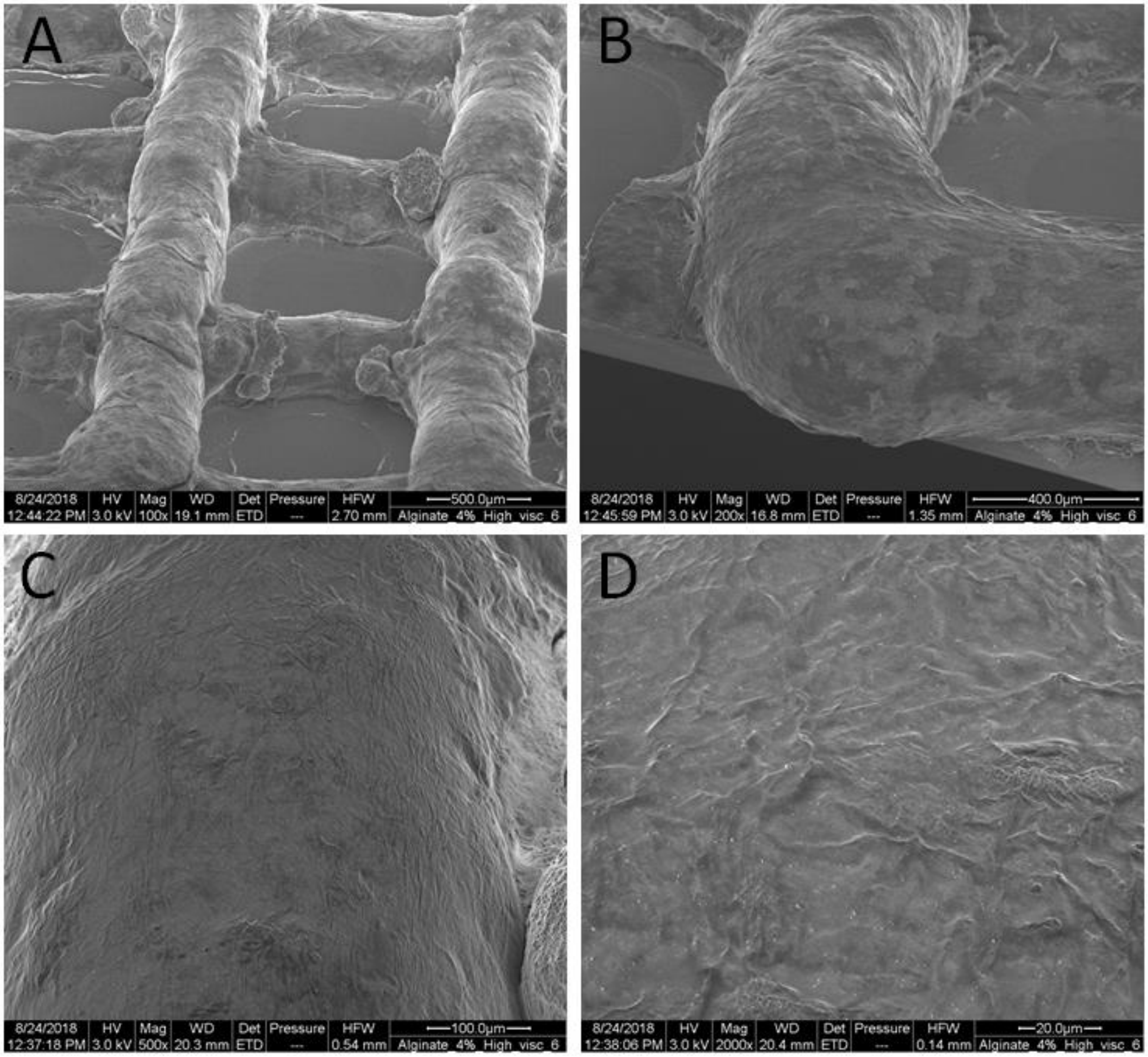


Figure S17. Morphological characterization of AlgH5 by SEM (A-D).

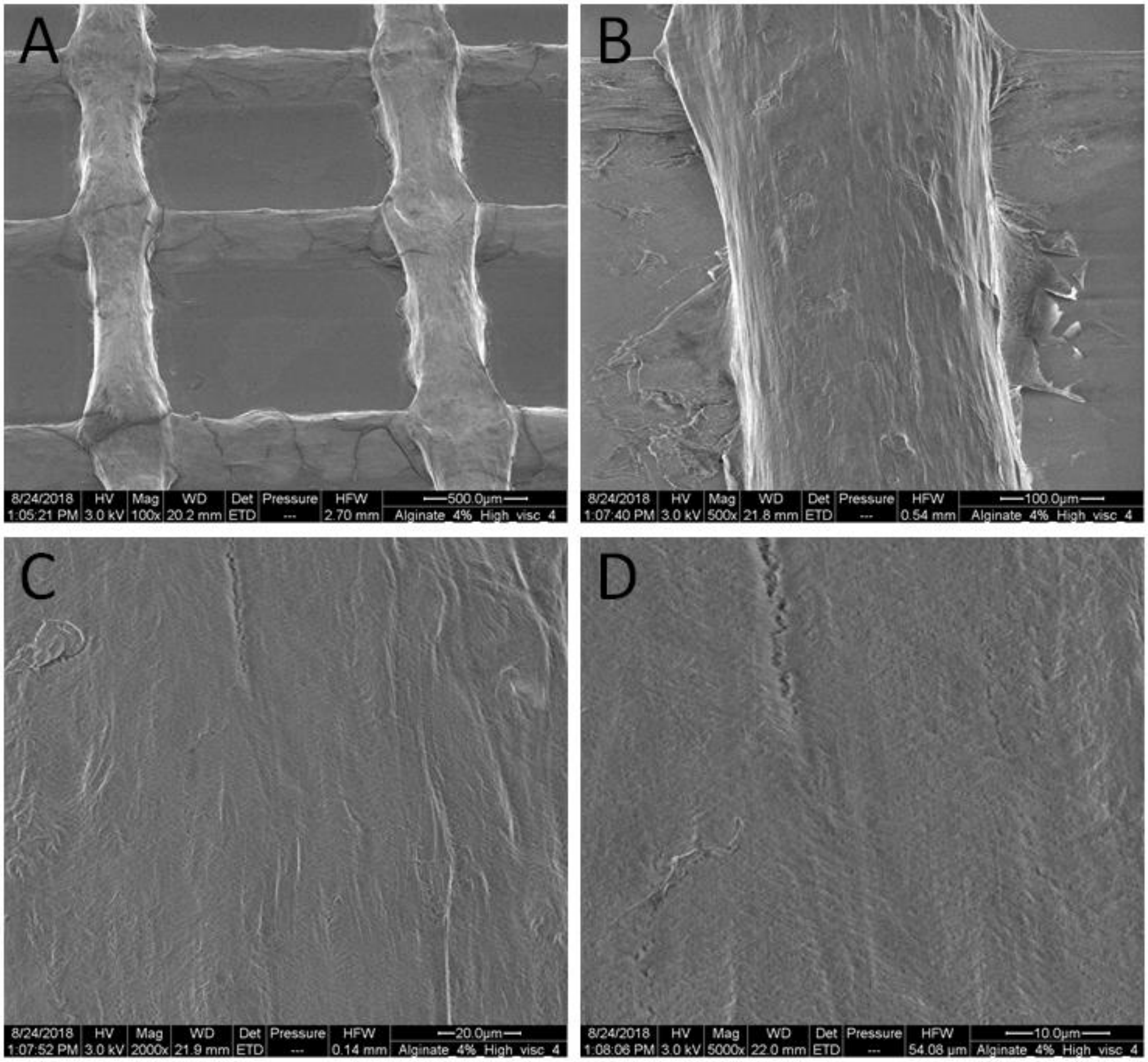


Figure S18. Morphological characterization of AlgH50 by SEM (A-D).
